# A computationally designed fluorescent biosensor for D-serine

**DOI:** 10.1101/2020.08.18.255380

**Authors:** Vanessa Vongsouthi, Jason H. Whitfield, Petr Unichenko, Joshua A. Mitchell, Björn Breithausen, Olga Khersonsky, Leon Kremers, Harald Janovjak, Hiromu Monai, Hajime Hirase, Sarel J. Fleishman, Christian Henneberger, Colin J. Jackson

## Abstract

Solute-binding proteins (SBPs) have evolved to balance the demands of ligand affinity, thermostability and conformational change to accomplish diverse functions in small molecule transport, sensing and chemotaxis. Although the ligand-induced conformational changes that occur in SBPs make them useful components in biosensors, they are challenging targets for protein engineering and design. Here we have engineered a D-alanine-specific SBP into a fluorescent biosensor with specificity for the signaling molecule D-serine (D-serFS). This was achieved through binding site and remote mutations that improved affinity (*K*_D_ = 6.7 ± 0.5 μM), specificity (40-fold increase *vs.* glycine), thermostability (Tm = 79 °C) and dynamic range (~14%). This sensor allowed measurement of physiologically relevant changes in D-serine concentration using two-photon excitation fluorescence microscopy in rat brain hippocampal slices. This work illustrates the functional trade-offs between protein dynamics, ligand affinity and thermostability, and how these must be balanced to achieve desirable activities in the engineering of complex, dynamic proteins.

## Introduction

Since the discovery of free D-serine in mammalian brain tissue in the early 1990s (Hashimoto et al., 1992), there has been considerable interest in its physiological and pathophysiological roles. It is now known that D-serine acts as a co-agonist of excitatory glutamate receptors of the N-methyl-D-aspartate receptor (NMDAR) subtype. Together with the primary agonist, L-glutamate, D-serine binds to the ‘glycine-binding site’ on the GluN1 ligand-binding domain (LBD) of the NMDAR to activate it (Panatier et al., 2006, Schell et al., 1995, Mothet et al., 2000). By the same mechanism, glycine also acts as a co-agonist of the NMDARs (Panatier et al., 2006, Schell et al., 1995, Mothet et al., 2000). Previous work has demonstrated the gating of synaptic NMDARs by D-serine (Henneberger et al., 2010, Papouin et al., 2012), and of extrasynaptic NMDARs by glycine (Papouin et al., 2012). Notably, it is synaptic NMDARs that are primarily responsible for inducing long-term potentiation (LTP) of excitatory synaptic transmission, a key mechanism of learning and memory (Bliss and Collingridge, 1993, Bliss and Cooke, 2011, Hardingham and Bading, 2010). D-serine has been associated with several conditions centered on cognitive impairment and disturbances to NMDAR activity. D-serine levels in the brain are largely regulated by the activity of serine racemase (SR), which racemizes L-serine to produce D-serine, and D-amino acid oxidase (DAAO), which catabolizes it (Wolosker et al., 1999, Pollegioni and Sacchi, 2010). Abnormal levels of both SR and DAAO have been associated with schizophrenia and Alzheimer’s disease (Basu et al., 2009, Labrie et al., 2009, Madeira et al., 2015). Recent studies in flies and mammals have also implicated D-serine in sleep regulation (Tomita et al., 2015, Liu et al., 2016, Papouin et al., 2017a, Dai et al., 2019) and kidney disease (Wiriyasermkul et al., 2020).

Despite the physiological importance of D-serine, aspects of D-serine signaling remain elusive. While it is thought that D-serine is a gliotransmitter released from astrocytes (Henneberger et al., 2010, Papouin et al., 2017b, Yang et al., 2003), its cellular origin continues to be an intensely debated topic (Wolosker et al., 2016, Papouin et al., 2017b). Progress towards understanding the D-serine pathway has been partly hindered by the lack of a suitable method to study the transmitter dynamically with high spatial and temporal resolution. While methods such as microdialysis can provide precise quantification of D-serine from tissues (Shippenberg and Thompson, 2001), they tend to suffer from poor temporal resolution and can cause mechanical damage to the tissue under study (Beyene et al., 2019, Ganesana et al., 2017). Amperometric probes for D-serine based on DAAO have also been developed by several groups (Pernot et al., 2008, Mohd Zain et al., 2012); however, these probes are subject to losses in sensitivity due to fouling of the outer biolayer and loss of enzymatic activity during operation (Dale et al., 2005), and also have a low spatial resolution.

Optical biosensors are an alternative to existing D-serine detection methods that have the potential to provide unparalleled spatial and temporal resolution when used with high resolution microscopy. Those based on Förster resonance energy transfer (FRET) are commonly a fusion of a ligand-binding domain to a pair of donor-acceptor fluorophores. Here, ligand-induced conformational changes in the binding domain lead to the displacement of the fluorophores and a measurable change in FRET efficiency (Kaczmarski et al., 2019). Several FRET-based biosensors of this architecture have been engineered for the dynamic study of key amino acids in brain tissue, including L-glutamate (Okumoto et al., 2005), glycine (Zhang et al., 2018), and L-arginine (Whitfield et al., 2015). The development of such a sensor for D-serine has previously been hindered by the absence of naturally occurring D-serine-specific solute-binding proteins (SBP).

Protein engineering can be used to overcome this issue and modify existing SBPs towards desirable binding affinity and specificity (Zhang et al., 2018). However, engineering complex dynamic molecules such as SBPs for affinity and specificity, while retaining high thermostability, can be challenging. In this study, we used rational and computational protein design to engineer a D-serine FRET sensor (D-serFS), using a D-alanine binding protein from *Salmonella enterica* (DalS) (Osborne et al., 2012) as a starting point. Iterative rounds of positive/negative design and experimental testing produced a robust, D-serine-specific fluorescent sensor with physiologically appropriate affinity for D-serine (*K*_D_ = 6.7 ± 0.5 μM) and reduced affinity for other physiologically relevant ligands, such as glycine. Additionally, the poor thermostability of the protein was improved through the addition of stabilizing mutations. *In-situ* testing of full-length D-serFS in acute hippocampal rat brain slices demonstrated that D-serFS was responsive to extracellular D-serine changes within a physiologically relevant concentration window.

## Results

### Homology-guided design of a D-serine-specific binding protein

An SBP from *Salmonella enterica*, DalS, that binds D-alanine and glycine (Osborne et al., 2012) was chosen as a starting point for the design of a D-serine-specific binding protein. The initial design was guided by a comparison to the GluN1 ligand-binding domain (LBD) of the NMDAR, a structural homolog of SBPs that contains the receptor’s D-serine/glycine-binding site (Furukawa and Gouaux, 2003). A structural alignment of DalS (PDB 4DZ1) and the GluN1 LBD (PDB 1PB8) (Fig. **1A**) revealed key differences in the polarity and cavity-size of the two binding sites, particularly at the residues comprising the D-alanine methyl side chain pocket: F117, A147 and Y148 (*vs.* L146, S180 and V181 in GluN1 LBD). Thus, these residues were targeted for mutagenesis towards D-serine specificity in DalS.

**Figure 1.**
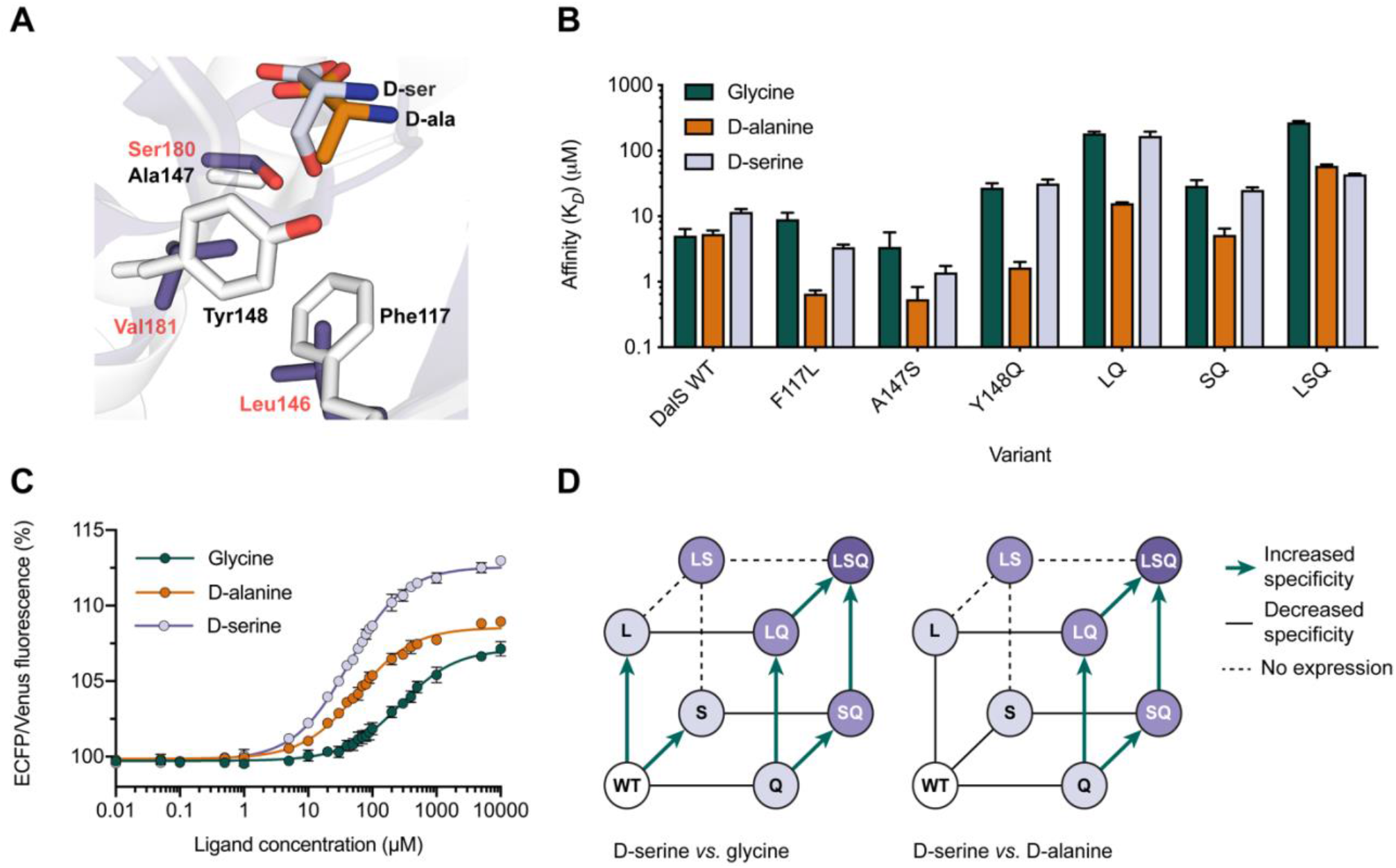
Homology-guided design of DalS. (A) Structural alignment of binding site residues targeted for mutagenesis in DalS (white, black labels; PDB 4DZ1) to homologous residues in the GluN1 LBD (purple, red labels; PDB 1PB8). (B) Binding affinities (μM) determined by fluorescence titration of wild-type DalS, and single, double and triple mutants, for glycine, D-alanine and D-serine (n = 3). (C) Sigmoidal dose response curves for F117L/A147S/Y148Q (LSQ) with glycine, D-alanine and D-serine (n = 3). Values are the (475 nm/530 nm) fluorescence ratio as a percentage of the same ratio for the apo sensor. (D) Schematic demonstrating the epistatic interactions between individual mutations contributing to D-serine specificity relative to glycine (left) and D-alanine (right). Increased D-serine specificity is represented by green arrows, whereas loss in specificity is represented by solid black lines. Dashed lines associated with the variant F117L/A147S (LS) represent the absence of experimental data for this variant due to no maturation of the Venus chromophore during expression.

Three mutations were initially considered: F117L, A147S and Y148Q. F117L was hypothesized to increase the size of the cavity and reduce van der Waals contacts with the methyl side chain of D-alanine, whilst A147S would increase the polarity of the binding site to form polar contacts with the hydroxyl side chain of D-serine. Y148 was initially mutated to glutamine rather than valine (as in the GluN1 LBD) to increase both the polarity and size of the side chain pocket relative to tyrosine, reducing potentially favorable interactions with D-alanine and glycine. Y148 was also mutated to valine (Y148V) in the background of a later variant of the sensor and this resulted in a large decrease in affinity for all ligands (*K*_D_ > 4000 μM) (SI. Fig. **1**).

To experimentally test the proposed design (F117L/A147S/Y148Q), DalS was first cloned between enhanced cyan fluorescent protein (ECFP) and Venus fluorescent protein (Venus); a commonly used FRET pair (Bajar et al., 2016). For immobilization in complex tissue, an N-terminal hexahistidine-tagged biotin domain was included in the construct. F117L, A147S and Y148Q were introduced to the wild-type FRET sensor in a combinatorial fashion. Soluble, full-length protein was obtained in high purity for all variants excluding the double mutant, F117L/A147S (LS). The LS mutant did not exhibit maturation of the Venus chromophore during expression, suggesting that the combination of the two mutations was too destabilizing to the binding protein that folding of the protein and the downstream Venus fluorescent protein were negatively affected.

Fluorescence titrations with D-alanine, glycine and D-serine were performed on the wild-type protein and the successfully purified variants to determine the binding affinities (*K*_D_) (Fig. **1B**, Table **1**). The wild-type exhibited similar affinity to D-alanine and glycine (~5 μM) and significant, but lower, affinity to D-serine (12 ± 1 μM). Among the single mutants, F117L and A147S increased affinity to D-alanine and D-serine by similar amounts, with minimal effect on glycine affinity. The only mutation that had a large effect in isolation was Y148Q, which had the unintended effect of increasing affinity to D-alanine and decreasing affinity ~8-fold to D-serine. In combination with Y148Q, F117L (LQ) decreased D-alanine affinity ~4.5-fold from Y148Q, while only decreasing D-serine affinity 3-fold, however, A147S (SQ), decreased D-alanine affinity ~3-fold, while increasing D-serine affinity ~3.5-fold. The effects of the mutations in the double mutants on glycine *vs.* D-serine were largely identical. Finally, the combination of the three mutations in F117L/A147S/Y148Q (LSQ), generated a variant that was specific for D-serine, exhibiting a *K*_D_ for D-serine of 43 ± 1 μM, compared to 59 ± 2 μM and 271 ± 16 μM for D-alanine and glycine, respectively (Fig. **1C**). The 6.3-fold higher affinity for D-serine *vs.* glycine is the more important comparison because of the spatial and temporal overlap between these in the brain; D-alanine, in contrast, is present in concentrations near or below detection levels in brain tissue (Hashimoto et al., 1992, Popiolek et al., 2018).

**Table 1.** Summary of the binding affinities for D-alanine, glycine and D-serine (μM), specificities and thermostabilities (°C) for all variants of the wild-type sensor. D-serine specificity of the variants relative to glycine (α_Gly_) and D-alanine (α_D-ala_) is defined as the ratio between the *K*_D_ for D-serine and that of the corresponding off-target ligand: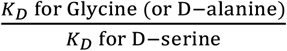. Values are mean ± s.e.m. throughout (n = 3).

| Variant | Binding Affinity, $K_D$ ($\mu\text{M}$) | | | Specificity | | $T_m$ ($^{\circ}\text{C}$) |
| --- | --- | --- | --- | --- | --- | --- |
| | D-ala | Gly | D-ser | $\alpha_{\text{Gly}}$ | $\alpha_{\text{D-ala}}$ | |
| WT | $5.4 \pm 0.4$ | $5 \pm 1$ | $12 \pm 1$ | 0.42 | 0.45 | $68.0 \pm 0.3$ |
| F117L | $0.65 \pm 0.03$ | $9 \pm 1$ | $3.7 \pm 0.2$ | 2.40 | 0.18 | $63.1 \pm 0.3$ |
| A147S | $0.62 \pm 0.03$ | $4.0 \pm 0.2$ | $1.8 \pm 0.4$ | 2.22 | 0.34 | $66.9 \pm 0.4$ |
| Y148Q | $1.6 \pm 0.3$ | $75 \pm 8$ | $101 \pm 3$ | 0.74 | 0.02 | $62.0 \pm 0.2$ |
| (LS) F117L/A147S<sup>†</sup> | - | - | - | - | - | - |
| (LQ) F117L/Y148Q | $32 \pm 2$ | $340 \pm 14$ | $309 \pm 19$ | 1.10 | 0.10 | $60.3 \pm 0.2$ |
| (SQ) A147S/Y148Q | $5.4 \pm 0.5$ | $29 \pm 2$ | $28 \pm 3$ | 1.04 | 0.19 | $60.8 \pm 0.2$ |
| (LSQ) F117L/A147S/Y148Q | $59 \pm 2$ | $271 \pm 16$ | $43 \pm 1$ | 6.30 | 1.44 | $60.5 \pm 0.2$ |
| (LSQ+E) LSQ + D216E | $118 \pm 3$ | $429 \pm 21$ | $41 \pm 1$ | 10.46 | 2.88 | $62.2 \pm 0.2$ |
| (LSQ+E+D) LSQ + A76D | $130 \pm 4$ | $410 \pm 100$ | $48 \pm 4$ | 8.54 | 2.71 | $65.0 \pm 0.1$ |
| (LSQED+K) LSQED + S60K | $180 \pm 1$ | $540 \pm 13$ | $61.0 \pm 0.2$ | 8.85 | 2.95 | $66.3 \pm 0.1$ |
| (LSQED+E) LSQED + S208E | $120 \pm 3$ | $420 \pm 93$ | $40 \pm 1$ | 10.50 | 3.00 | $66.2 \pm 0.1$ |
| (LSQED+K) LSQED + N200K | $230 \pm 63$ | $1400 \pm 1000$ | $470 \pm 270$ | 2.98 | 0.49 | $69.4 \pm 0.1$ |
| (LSQED+D) LSQED + T172D | - | - | - | - | - | $71.0 \pm 0.7$ |
| (LSQED+R) LSQED + S60R | $202 \pm 42$ | $1200 \pm 380$ | $70 \pm 17$ | 17.14 | 2.89 | $79.0 \pm 0.3$ |
| (LSQED+Y) LSQED + T197Y | $13 \pm 1$ | $41 \pm 17$ | $7.0 \pm 0.4$ | 5.86 | 1.86 | $79.0 \pm 0.3$ |
| (LSQED+RY) LSQED + S60R/T197Y | $14 \pm 1$ | $49 \pm 10$ | $6.1 \pm 0.5$ | 8.03 | 2.30 | * |
| **Full-length LSQED+Y (D-serFS)** | $14 \pm 1$ | $40 \pm 2$ | $4.8 \pm 0.2$ | 8.33 | 2.92 | $78.8 \pm 1.0$ |

<sup>†</sup> This variant did not yield mature fluorescent protein.
\* Indicates that a sigmoidal transition in fluorescence ratio as a function of increasing temperature was not observed within the temperature range of the experiment (20 – 85 $^{\circ}\text{C}$ ), suggesting a $T_m$ higher than $\sim 85$ $^{\circ}\text{C}$ .

The combinatorial analysis of the effects of these mutations is indicative of strong epistasis, i.e. the effects of the mutations are not additive and there is no smooth pathway of stepwise increases in D-serine specificity available by which we could arrive at the LSQ variant (Fig. **1D**). This strong epistasis highlighted the advantage of adopting a rational design approach to engineering the binding site: rational design allows for the introduction of all three mutations simultaneously, whereas proceeding via single mutational steps would have resulted in dysfunctional “dead end” single or double mutants.

### Increasing D-serine specificity of LSQ using ligand docking

In order to identify mutations to the binding site that would further increase the specificity of LSQ for D-serine, modelling of the LSQ binding protein with FoldX (Schymkowitz et al., 2005) and ligand docking with Glide (Friesner et al., 2004), were performed. As LSQ maintained similar affinities for D-alanine (*K*_D_ = 59 ± 2 μM) and D-serine (*K*_D_ = 43 ± 1 μM), both ligands were docked into the binding site of a model of LSQ and the top poses were analyzed (Fig. **2A**). In agreement with previous work (Osborne et al., 2012), the poses suggested that R102 and S97 stabilize the carboxylate group of all docked ligands. It appears that D216, N115, and Y173, likely play roles in ligand binding and specificity given their proximity to the side chains. Thus, these residues were rationally mutated *in silico* for subsequent rounds of ligand docking.

**Figure 2.**
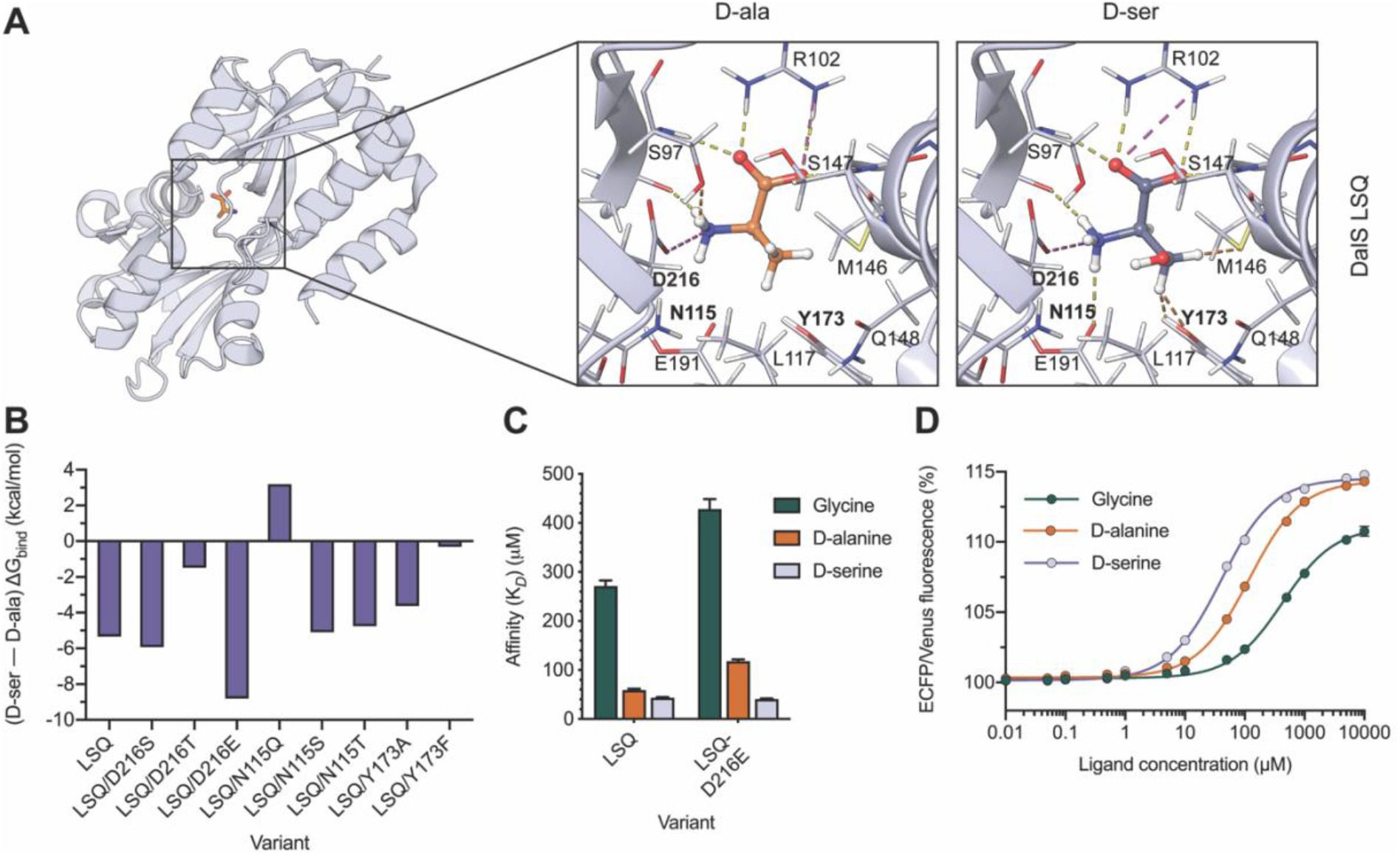
Ligand docking of the LSQ binding protein using Glide (Friesner et al., 2004). (A) Top binding poses of D-alanine (orange, left) and D-serine (purple, right) in a FoldX-generated (Schymkowitz et al., 2005) model of DalS with mutations F117L, A147S, and Y148Q (DalS LSQ). Dashed lines represent non-covalent interactions between binding site residues and the ligands, including hydrogen bonds (yellow), salt-bridges (pink) and unfavourable clashes (red). Residues targeted for further rational design, N115, Y173 and D216, are bolded. (B) Difference between ΔG_bind_ (kcal/mol) for D-serine and D-alanine, predicted using Prime MM-GBSA (Rapp et al., 2011) for FoldX-generated models of DalS LSQ and single mutants in the background of DalS LSQ. (C) Binding affinities (μM) determined by fluorescence titration of LSQ and LSQ/D216E (LSQE), for glycine, D-alanine and D-serine (n = 3). (D) Sigmoidal dose response curves for LSQE with glycine, D-alanine and D-serine (n = 3). Values are the (475 nm/530 nm) fluorescence ratio as a percentage of the same ratio for the apo sensor.

We first mutated each residue to the corresponding residue in the GluN1 LBD, giving rise to LSQ-D216S,-N115Q, and -Y173A. As the carboxylic acid of D216 was predicted to form stabilizing interactions with the amine of both D-alanine and D-serine, the mutation of this residue to glutamate, D216E, was performed. The introduction of other polar residues at positions 115 and 216 was also explored, leading to the mutants LSQ-D216T,-N115T, and -N115S. Lastly, as the side chain of Y173 was predicted to clash with the D-serine ligand in LSQ, the mutation of this residue to phenylalanine, Y173F, was investigated. D-serine and D-alanine were docked into models of the proposed mutants of LSQ and the generated poses were analyzed by visual inspection. In order to quantify and compare the effects of the mutations on binding specificity, Prime MM-GBSA was used to predict the free energy of binding (ΔG_bind_) in kcal/mol for each ligand (SI. Table **1**). Prime MM-GBSA can effectively predict the relative, rather than absolute, binding free energies for congeneric ligands (Lyne et al., 2006, Rapp et al., 2011). Due to this, focus was placed on the difference between the ΔG_bind_ for D-serine and D-alanine for each mutant, where positive and negative values corresponded to preferential binding of D-alanine and D-serine, respectively (Fig. **2B**). Using this approach, D216E appeared to be the most beneficial mutation for increasing D-serine specificity (ΔG_bind_ difference = −8.8 kcal/mol) relative to LSQ (ΔG_bind_ difference = −5.3 kcal/mol) and was selected for experimental characterisation. The remaining single mutations did not exhibit ΔG_bind_ differences that were indicative of increased D-serine specificity.

Experimental testing of LSQ/D216E (LSQE) revealed improved specificity towards D-serine. Fluorescence titrations showed that it decreased the affinity of the sensor for both D-alanine (*K*_D_ = 118 ± 3 *vs.* 59 ± 2 μM) and glycine (*K*_D_ = 429 ± 21 *vs.* 271 ± 16 μM), whilst the affinity for D-serine was slightly increased (*K*_D_ = 41 ± 1 *vs.* 43 ± 1 μM) compared to LSQ (Fig. **2C – D**). Furthermore, this improvement in specificity did not affect the dynamic range of the sensor. As a result, LSQE was taken forward for an additional round of engineering.

### Stabilizing mutations in LSQE improve the affinity and specificity for D-serine

Preliminary testing of the LSQ variant under two-photon excitation (2PE) fluorescence microscopy showed a decrease in the dynamic range following the reconstitution of lyophilised sensor (SI. Fig. **2**). As ECFP/Venus is a common FRET pair, and we had successfully used this sensor construct in previous sensors that had not suffered a loss in dynamic range following this process (Zhang et al., 2018, Whitfield et al., 2015), this effect was attributed to the inadequate stability of the solute binding protein. The thermostability of each sensor variant was determined by measuring the fluorescence ratio (ECFP/Venus) as a function of increasing temperature (Fig. **3A**). This analysis revealed that all of the variants were less thermostable than the wild-type (by up to 8 °C), which we assume is very close to the point at which the protein can no longer fold, as was observed for the F117L/A147S variant. The destabilizing effects of these mutations are consistent with the observation that mutations to active/binding/core sites are more often than not destabilizing (Tokuriki et al., 2007).

**Figure 3.**
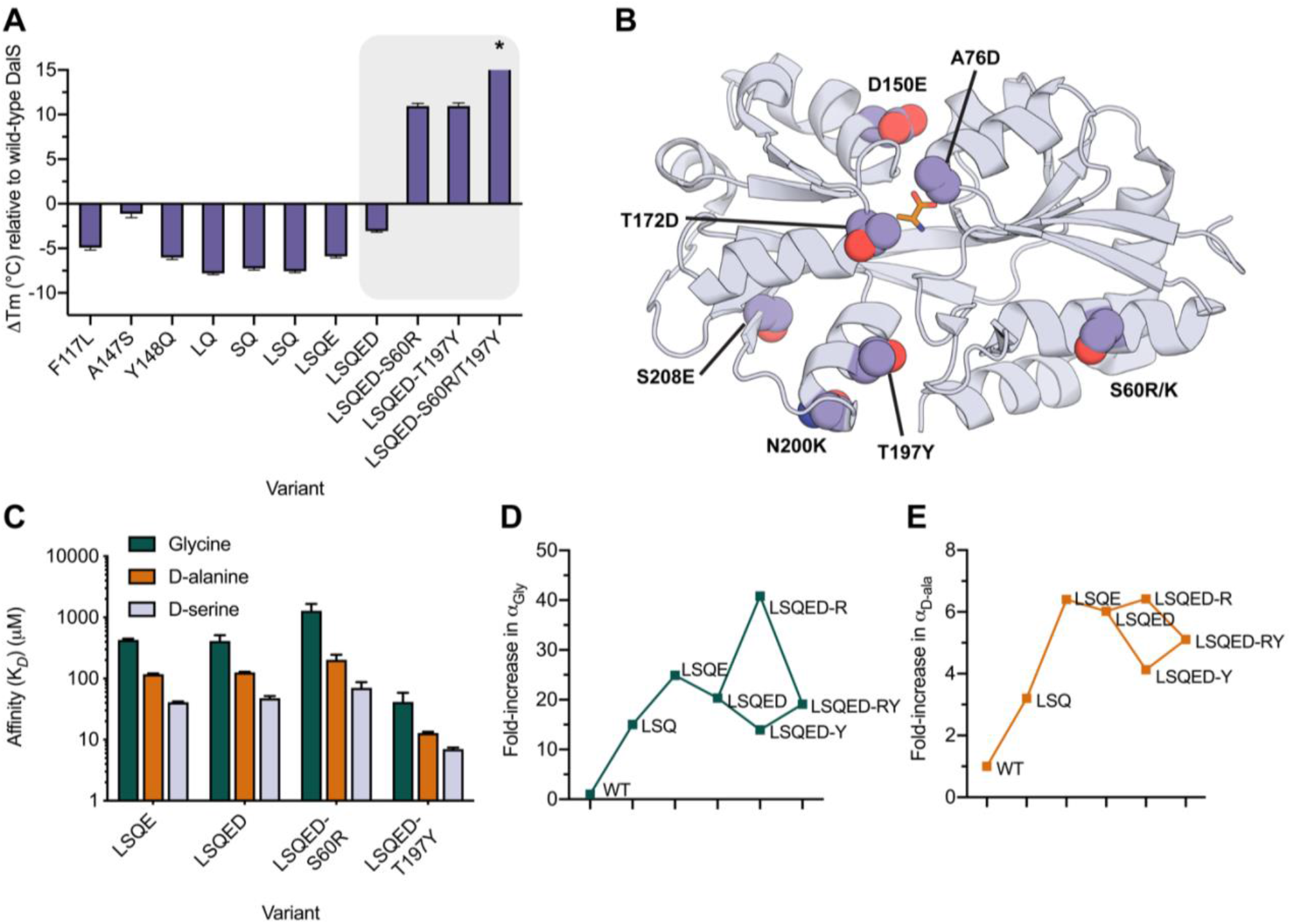
(A) Change in melting temperature of DalS variants relative to the wild-type (ΔTm) (°C) (n = 3). Mutations to the binding site are destabilising. Stabilising mutations to LSQE predicted by FoldX (Schymkowitz et al., 2005) and PROSS (Goldenzweig et al., 2016) are highlighted (grey box). *The ΔTm for LSQED-S60R/T197Y exists outside the axis limit as the unfolding of this variant could not be detected within the temperature range of the experiment (20 – 85 °C). (B) Crystal structure of wild-type DalS (PDB 4DZ1) with the positions of predicted stabilising mutations highlighted (spheres) and labelled. (C) Binding affinities (μM) determined by fluorescence titration of LSQE, LSQE-A76D (LSQED), LSQED-S60R (LSQED-R), and LSQED-T197Y (LSQED-Y), for glycine, D-alanine and D-serine (n = 3). In (D) – (E), the fold-increase in D-serine specificity compared to the wild-type for key variants in the design of D-SerFS, relative to glycine (D) and D-alanine (E) is shown. Specificity (α) is defined as: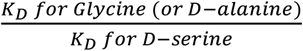.

In order to identify potentially stabilizing mutations in the binding protein, two computational methods were used: the structure-based FoldX (Schymkowitz et al., 2005) and the sequence and structure-based PROSS (Goldenzweig et al., 2016). Residues on the surface (S60, A76, D150, T172, N200, S208; Fig. **3B**) were initially targeted for mutation by FoldX to reduce the likelihood of destabilizing mutations (Tokuriki et al., 2007). The mutations to each of the other amino acids at each position were ranked by the predicted change in Gibbs free energy of folding (ΔΔG), and the most stabilizing mutation at each position that introduced a polar or charged residue to the surface was identified. This produced the following set of mutations: A76D, S208E, S60K, T172D, S60R, N200K, and D150E (Fig. **3B**,Table **1**). In parallel, the PROSS webserver developed by Goldenzweig et al. (Goldenzweig et al., 2016) was used to identify stabilizing mutations. The PROSS algorithm combines phylogenetic analysis and Rosetta-based design to provide users with combinatorial designs predicted to have improved stability relative to the input sequence and structure. A PROSS calculation was performed on the DalS crystal structure and residues with known importance to ligand binding were not allowed to design. This produced the following set of mutations: N32R, S60K, S61A, Q64K, S65E, H67G, L70C, T73V, A76S, G82A, Q88K, E110D, I114T, A123K, H125D, N131D, N133S, N136K, T176A, T176V, T197Y, K199L, L234D, Q242E, S245K, S245Q, G246A, G246E. The individual mutations identified by PROSS were also evaluated with FoldX. All the identified stabilizing mutations were then ranked by the ΔΔG values calculated by FoldX (SI. Table **2**).

Initially, the A76D mutation, which was predicted by FoldX to be the most stabilizing, was selected for experimental testing in the background of the LSQE variant. LSQE/A76D (LSQED) displayed a +3 °C improvement in thermostability (Tm = 65.0 ± 0.1 °C) compared to LSQE and resulted in little change to the binding affinities (Fig. **3C**, Table **1**). Given this improvement, all mutations with a FoldX ΔΔG that was more negative than 2.5 FoldX standard deviations (< −1.15 kcal/mol) were individually introduced to the background of LSQED. Two variants, LSQED/S208E and LSQED/S60K, displayed no significant improvements (< 1 °C) in thermostability (Table 1). The variants, LSQED/T172D and LSQED/N200K exhibited moderate improvements in thermostability (Tm = 71.0 ± 0.7 °C and 69.4 ± 0.1 °C, respectively), while the variants LSQED/S60R (LSQED-R) (Tm = 79.0 ± 0.3 °C) and LSQED/T197Y (LSQED-Y) (79 ± 0.3 °C) resulted in the greatest improvements in thermostability (Fig. **3A**, Table **1**).

Unexpectedly, several of the mutations that were identified to increase thermostability were found to have pronounced effects on substrate affinity and specificity. For example, N200K resulted in a ~10-fold reduction in affinity for D-serine (Table **1**). In contrast, S60R significantly decreased the affinity for glycine (*K*_D_ = 1200 ± 380 μM; Fig **3C**). In a comparison of the fold-change in specificity of key variants relative to the wild-type (Fig. **3D – E**), S60R was found to be the most specific variant, exhibiting a 40-fold and 6-fold greater specificity for D-serine, relative to glycine and D-alanine, respectively, albeit with slightly reduced affinity for D-serine (*K*_D_ = 70 ± 17 μM; Fig. **3C**). The T197Y mutation was also unexpectedly beneficial for binding and resulted in a marked increase in affinity for D-serine (*K*_D_ = 7.0 ± 0.4 μM; Fig. **3C**), as well as for the native ligands D-alanine (*K*_D_ = 13 ± 1 μM) and glycine (*K*_D_ = 41 ± 17 μM). The T197Y mutant, LSQED-Y, maintained D-serine specificity, displaying a 14-fold and 4-fold greater specificity for D-serine, relative to glycine and D-alanine, respectively, compared to the wild-type (Fig. **3D – E**). Interestingly, when the mutations S60R and T197Y were included together in the same variant (LSQED-RY), the affinities for D-serine (*K*_D_ = 6.1 ± 0.5 μM), D-alanine (*K*_D_ = 14 ± 1 μM) and glycine (*K*_D_ = 49 ± 10 μM) closely resembled that observed for the T197Y mutation alone (**Table 1**). As no sigmoidal transition in fluorescence ratio as a function of increasing temperature was observed for this variant, this suggested that the thermostability is similar to, or exceeds, the upper limit of the temperature range of the experiment (~ 85 °C; Fig. **3A**).

### The molecular basis for the effects of remote mutations on ligand affinity

The large effect of the T197Y mutation on improving the binding affinities to the three ligands, despite being ~13 Å from the binding site, was unexpected, prompting further analysis of the molecular basis for this effect. There is no apparent structural explanation for why this remote mutation would change affinity based on the crystal structure of DalS, i.e. there is no mechanism by which it could change the shape or the protein-ligand interactions of the binding site. As the crystal structure of DalS (PDB 4DZ1) is in the ligand bound (closed) conformation, and it is known this protein family fluctuates between open and closed states, we investigated the open-closed conformational transition of the protein scaffold to examine whether the mutation was altering affinity by differentially stabilizing the ligand-bound (closed) state over the unbound (open) state. To do this, we first needed a model of the unbound (open) state. Previous work on SBPs closely related to DalS have shown that molecular dynamics (MD) simulations of closed SBP crystal structures in the apo state can reasonably sample open conformational substates (Clifton et al., 2018, Kaczmarski et al., 2020). To simulate the closed to open conformational transition of DalS (PDB 4DZ1), MD simulations (100 ns x 10 replicates) were run with the ligand removed, and clustering analysis was performed to obtain a representative open conformation from the largest cluster. The Cα–Cα distance between residues A85 and K153 (either side of the binding site cleft), and radius of gyration, were calculated for all frames of the simulation (Fig. **4A**). The residue 85-153 Cα–Cα distance in the open conformation was 0.9 nm greater than that observed in the closed crystal structure (3.0 *vs*. 2.1 nm; Fig. **4B**). In the X-ray crystal structures of a closely related SBP, AncCDT-1, the difference in Cα–Cα distance (at equivalent positions) between the open (PDB 5TUJ) and closed (PDB 5T0W) crystallographic states is similar (0.8 nm) (Clifton et al., 2018, Kaczmarski et al., 2020). The difference in the radius of gyration (~1 Å) between the open and closed crystallographic states of AncCDT-1 was also comparable to that observed between the representative open conformation and the closed crystal structure of DalS.

**Figure 4.**
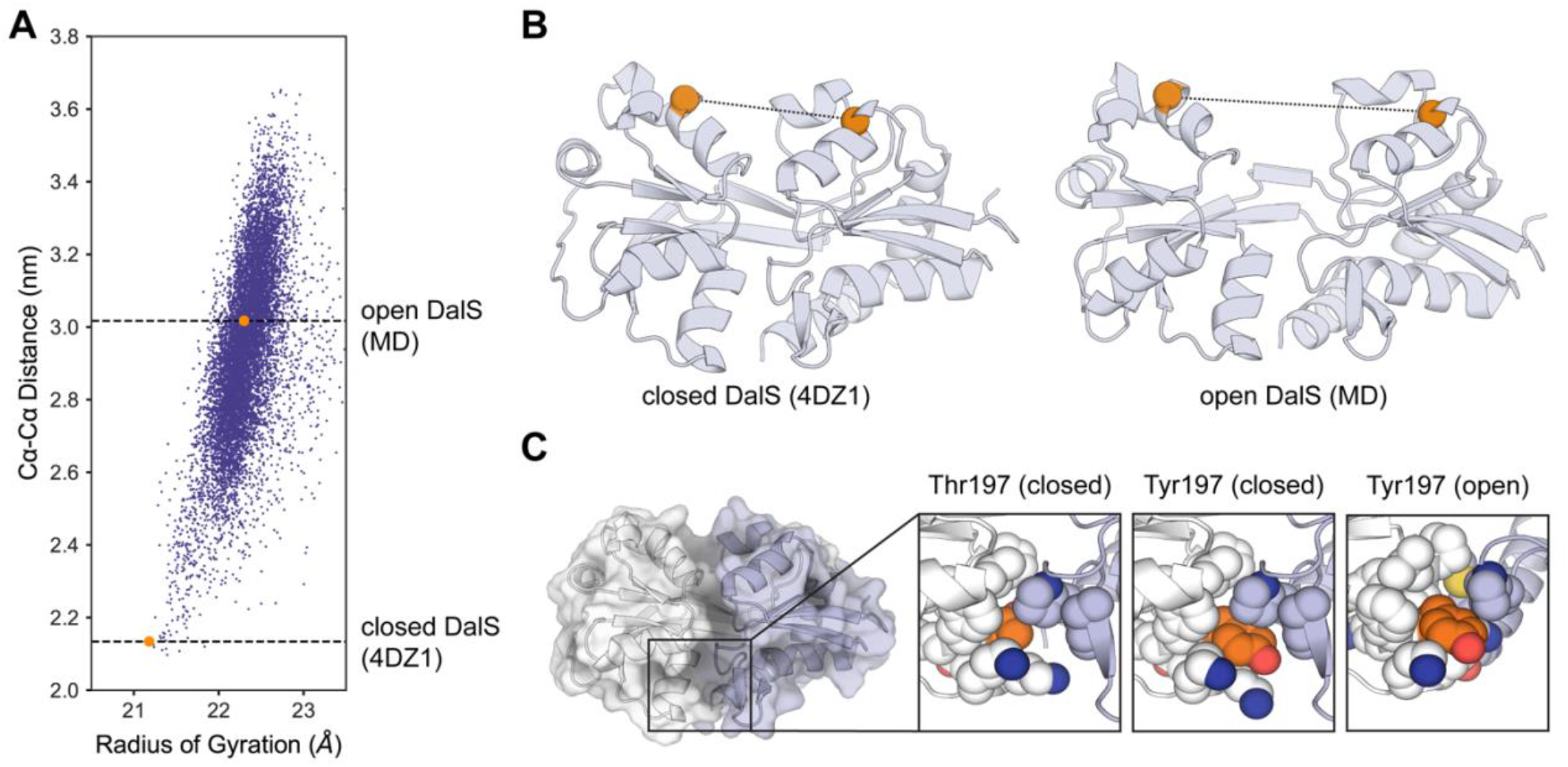
(A) Conformational substates of DalS sampled by MD simulations. The Cα–Cα distance between A85 and K153 is plotted against the radius of gyration where each data point represents a single frame of the MD simulation (sampled every 0.1 ns). Points corresponding to the closed crystal structure of DalS (PDB 4DZ1) and the representative open conformation are highlighted in orange. (B) The structure of closed DalS (left) is compared to the structure of the representative open conformation. Residues A85 and K153 are represented by orange spheres, highlighting the difference in Cα–Cα distance. (C) Thr197 in the closed crystal structure is compared to FoldX-generated (Schymkowitz et al., 2005) models of the T197Y mutation in the open and closed conformations. Residue 197 (orange spheres) and residues within 4 Å (white spheres) are displayed, showing tighter packing of Tyr197 by surrounding residues in the closed conformation compared to the open conformation. The positioning of Tyr197 at the interface between Lobe 1 (purple) and Lobe 2 (white) is also highlighted.

T197Y was modelled into the representative open conformation and the closed crystal structure of DalS using FoldX (Schymkowitz et al., 2005). A comparison of Thr and Tyr at position 197 in the closed conformation (Fig. **4C**) showed that the mutation to the larger residue, Tyr, fills a void in the protein between the two lobes and is likely to increase stability by improving hydrophobic packing. Given that residue 197 is positioned at the interface between the two lobes of the binding protein, the improved hydrophobic packing upon mutation to Tyr will stabilize the interaction between them and is likely to stabilize the closed state (relative to Thr). Contrastingly, in the open conformation (Fig. **4C**), Tyr197 is less tightly packed by surrounding residues and is more solvent exposed. This is consistent with the predicted effects on stability as calculated by FoldX, with the calculated ΔΔG for introducing the T197Y mutation to the open and closed conformations being −0.7 and −1.6 kcal/mol, respectively. This was also investigated experimentally: the fluorescence spectra of the LSQED-Y mutant, which displayed a reduced dynamic range *in-vitro* (7%), is consistent with a reduced population of the open state (SI. Fig. **3**). These results suggest that the remote mutation affects affinity by stabilizing the ligand bound state, which is also consistent with the observation that the affinity increases for all ligands rather than specifically for D-serine (Table **1**), providing a molecular-level explanation for the unexpected effect of this remote mutation on ligand affinity.

### Extending a truncation in D-serFS for improved dynamic range

The LSQED-Y variant, hereafter referred to as D-serFS (D-serine FRET Sensor), exhibited appropriate binding affinities for the physiological detection of D-serine. However, the relatively low dynamic range of this variant (7%) limited its use. Sequencing of the full D-serFS construct revealed an unintentional truncation of 10 amino acids at the C-terminus of Venus and a 6 amino acid insertion from the vector backbone to the C-terminus of Venus (Fig. **6A**). Full length D-serFS was re-cloned as a sensor construct, expressed and purified for characterization. Full-length D-serFS displayed an improved dynamic range (14.7 ± 0.1%) in response to D-serine *in-vitro* (Fig. **6B**) while maintaining similar binding affinity (*K*_D_ for D-serine = 4.8 ± 0.2 μM) (Fig. **6C**, SI. Fig. **4**) and thermostability (Tm = 78.8 ± 1.0 °C) (Table **1**) compared to the truncated variant. Thus, while the truncated version was suitable for the engineering work to optimize the affinity, thermostability and specificity of the binding domain, Full-length D-serFS was used for subsequent sensing applications.

**Figure 5.**
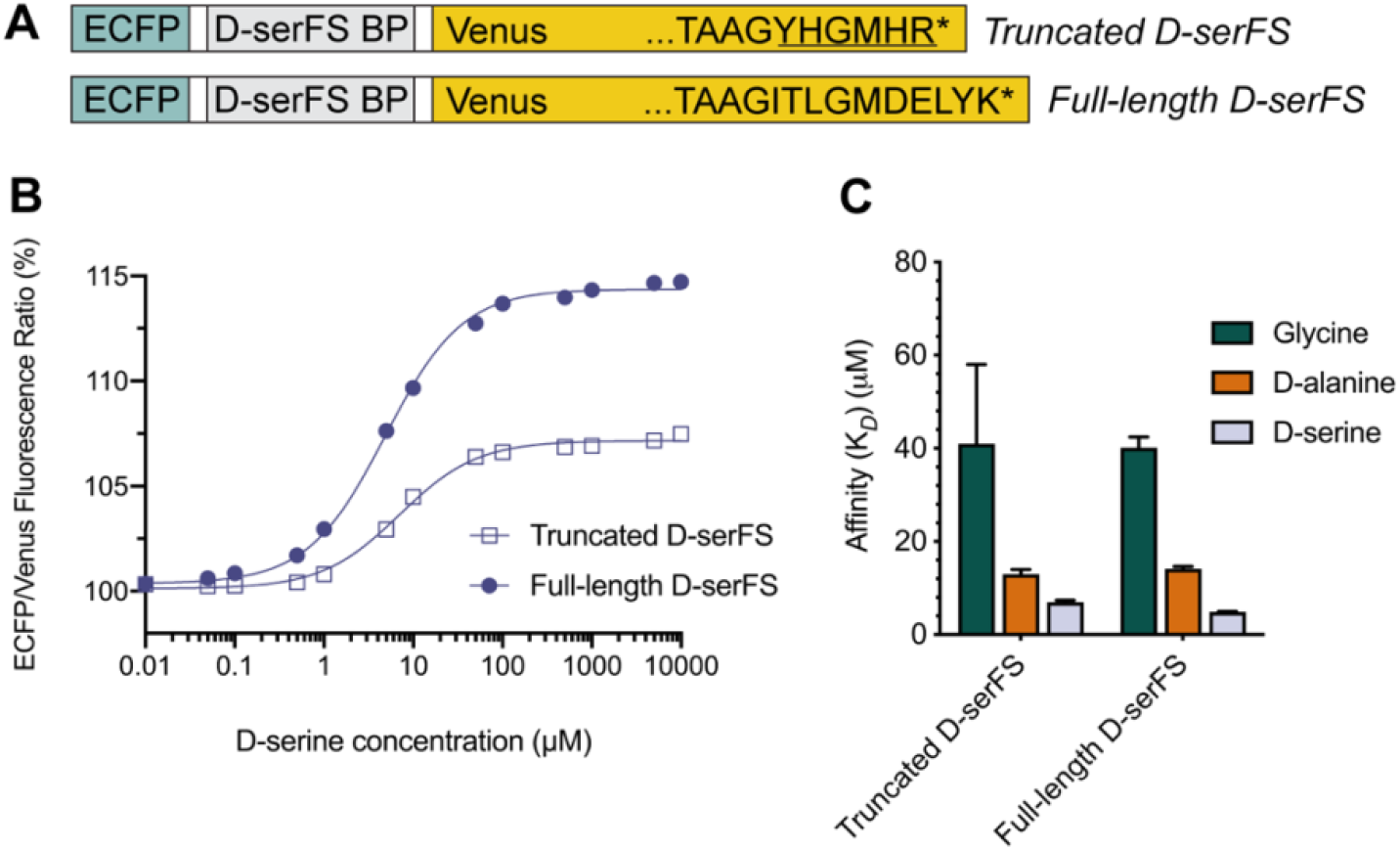
Characterization of full-length D-serFS. (A) Schematic showing the ECFP (blue), D-serFS binding protein (D-serFS BP; grey) and Venus (yellow) domains in D-serFS. The C-terminal residues of the Venus fluorescent protein sequence are labelled, showing the truncated (top) and full-length (bottom) C-terminal sequences. The underlined amino acids in truncated D-serFS represent residues introduced from the backbone vector sequence during cloning. *Represents the STOP codon. (B) Sigmoidal dose response curves for truncated and full-length D-serFS with D-serine (n = 3). Values are the (475 nm/530 nm) fluorescence ratio as a percentage of the same ratio for the apo sensor. (C) Binding affinities (μM) determined by fluorescence titration of truncated and full-length D-serFS, for glycine, D-alanine and D-serine (n = 3).

**Figure 6.**
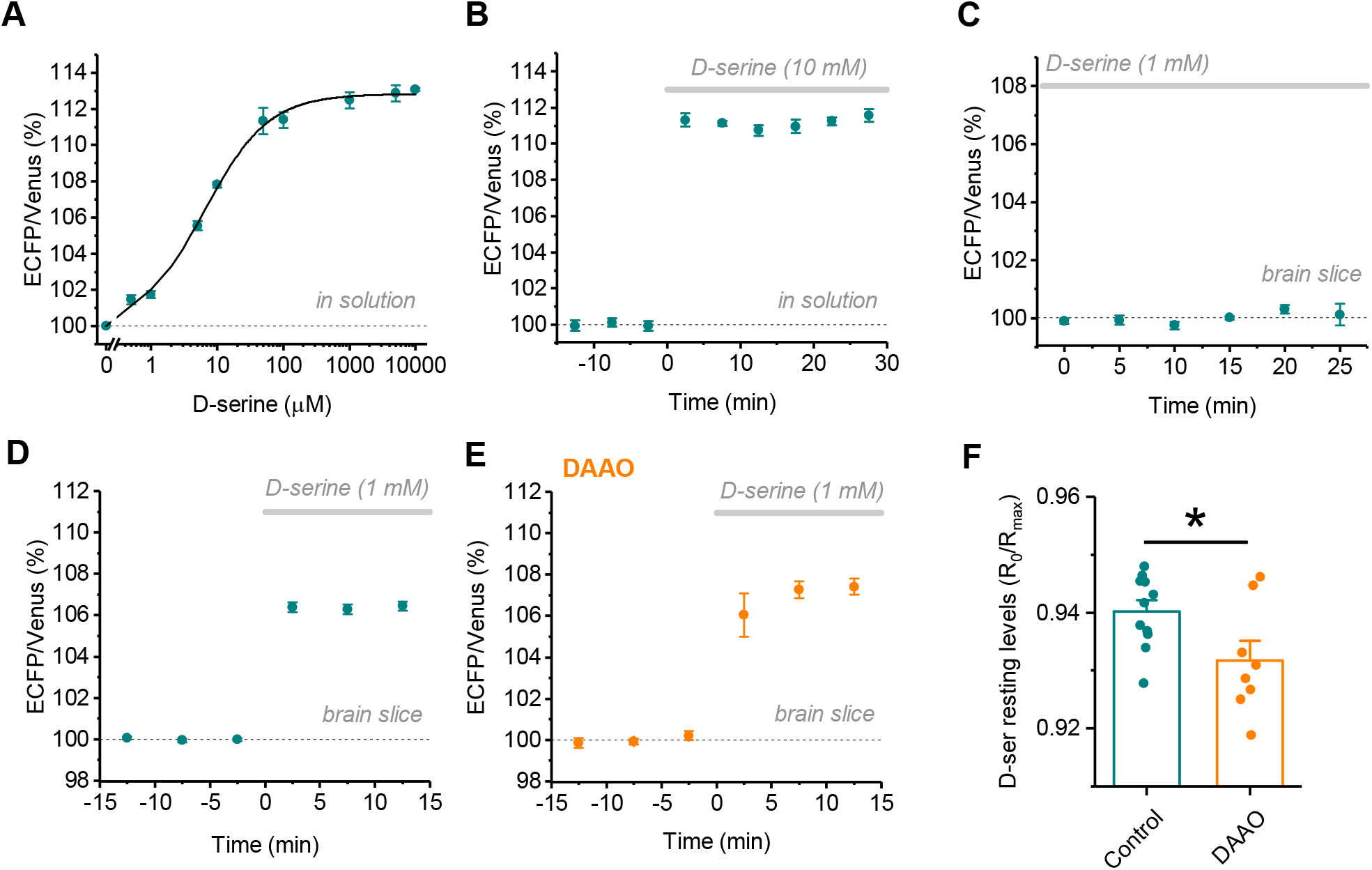
Characterization and in-situ testing of full-length D-serFS using 2PE fluorescence microscopy. (A) Calibration curve of full-length D-serFS using 2PE (λ_exc_ = 800 nm) shows a dose-dependent change in the ECFP/Venus fluorescence intensity ratio. Fitting the relationship between fluorescence ratio and D-serine concentration with a Hill equation obtains a *K*_D_ of 6.7 ± 0.5 μM (dynamic range 12.8 ± 0.2 %, n = 3 independent experiments). (B) Monitoring of the ECFP/Venus ratio in a meniscus after applying 10 mM D-serine demonstrates a stable fluorescence ratio over a time course of 30 minutes (n = 5). (C) Monitoring of the ECFP/Venus ratio in rat hippocampal slices during persistent exposure of the sensor to 1 mM D-serine demonstrates a stable fluorescence ratio also at this D-serine concentration and in the organized brain tissue over a time course of 25 minutes (n = 4). (D) Imaging of full-length D-serFS in response to the exogenous application of 1 mM D-serine in rat brain hippocampal slices (n = 11). (E) Imaging of full-length D-serFS in response to the exogenous application of 1 mM D-serine in rat hippocampal slices incubated with DAAO (n = 8). (F) To estimate the resting concentration of D-serine in control slices (n = 11) and slices incubated with DAAO (n = 8), ECFP/Venus fluorescence intensity ratios R_0_ and R_MAX_ were calculated, where R_0_ and R_MAX_ represent the resting ratio and the ratio in the presence of a saturating D-serine concentration of 1 mM (p = 0.03, two-sided Student’s t-test). Error bars represent ± s.e.m. * p < 0.05.

### Performance of full-length D-serFS in hippocampal tissue

The full-length D-serFS variant was used for *in-situ* testing as a biosensor due to its high D-serine affinity, specificity for D-serine over glycine, and thermostability. Furthermore, additional fluorescence titrations of full-length D-serFS *in-vitro* confirmed that the sensor did not bind other small molecules prominent in brain tissue (L-serine, GABA, L-glutamate, L-aspartate) with any significance (SI. Fig. **4**). To test the compatibility of full-length D-serFS with 2PE fluorescence microscopy, the sensor was imaged at an excitation wavelength of 800 nm in free solution under the objective and titrated with increasing concentrations of D-serine. In these conditions, the sensor exhibited similar affinity and dynamic range to that observed *in vitro* previously (Fig. **6A**), confirming that full-length D-serFS would be suitable for use with 2PE and thus, within intact tissue. Imaging the sensor over time in a meniscus after applying a saturating concentration of D-serine (10 mM) demonstrated that the ECFP/Venus ratio remained stable over a time course of at least 30 minutes (Fig. **6B**).

Next, full-length D-serFS was tested in 350 μm thick acute slices of rat hippocampus. The sensor was anchored in the extracellular space of the CA1 region of biotinylated acute hippocampal slices using a biotin-streptavidin linker, as described previously. The ECFP/Venus ratio did not change in the presence of saturating D-serine concentrations for up to 25 minutes (Fig. **6C**), indicating that the senor is stable in this environment. Furthermore, a rapid change in the ECFP/Venus fluorescence ratio was observed following the bath application of 1 mM D-serine to the hippocampal slice (Fig. **6D**). To further test if full-length D-serFS is able to report changes in extracellular D-serine levels, we recorded ECFP/Venus fluorescence following the bath application of 1 mM D-serine in brain slices incubated with the D-serine degrading enzyme DAAO (Fig. **6E**). Similar to previous experiments (Zhang et al., 2018), we used the change of the ECFP/Venus fluorescence intensity ratio (R) under baseline conditions (R_0_) relative to the ratio in the presence of saturating extracellular D-serine concentrations (R_MAX_) as a measure of D-serine resting levels. We found that D-serFS in slices incubated with DAAO reported significantly reduced resting levels of D-serine compared to the control slices (Fig. **6F**). It should be noted that it is not known how efficiently DAAO degrades extracellular D-serine in brain slices and to what exact value extracellular D-serine concentrations are decreased. Irrespective of that, these experiments qualitatively demonstrate that D-serFS can report changes of ambient extracellular D-serine levels in brain tissue.

## Discussion

In this study, we have constructed the optical sensor D-serFS and demonstrated its utility in detecting D-serine, an NMDAR co-agonist. In the absence of a suitable D-serine binding protein to use as a recognition domain for the sensor, we used rational and computational design to alter the specificity of an existing D-alanine/glycine binding protein from *Salmonella enterica* (DalS) (Osborne et al., 2012). An initial round of rational design guided by homology to the GluN1 ligand-binding domain of the NMDAR (Furukawa and Gouaux, 2003), followed by FoldX (Schymkowitz et al., 2005) modelling and Glide (Friesner et al., 2004) ligand docking, led to a variant of DalS (LSQE) that displayed D-serine binding specificity (*vs*. other amino acids). The stepwise introduction of individual binding site mutations towards the proposed rational design demonstrated that there was no single mutational pathway to the improved LSQ variant that displayed improvements at each step owing to the epistatic interactions between these residues. Thus, it was only by introducing all three mutations simultaneously (F117L, A147S, Y148Q) that the improvements to D-serine specificity could be achieved, highlighting the advantage of rational computational design over evolutionary approaches in some instances. This allowed us to achieve the improvement in specificity through testing a small number of variants rather than large libraries. However, the affinity of this first round variant (LSQ) for D-serine (*K*_D_ = 41 ± 1 μM) was too low to be useful in some biological contexts, such as in neuroscience. Measurement of the thermostabilities of the mutated variants revealed an overall decrease in thermostability with increased D-serine specificity. As the sensor was required to be robust against the effects of lyophilization and reconstitution, as well as stable in the crowded environment of brain tissue, we employed the computational methods FoldX (Schymkowitz et al., 2005) and PROSS (Goldenzweig et al., 2016) to identify stabilizing mutations to the binding protein. As the mutations predicted by FoldX and the Rosetta-ddG application have been shown to overlap in only 12 – 25% of all predictions (Buß et al., 2018, Wijma et al., 2014), combining both algorithms had the potential to provide greater coverage of beneficial mutations. In the background of LSQE, the combination of a stabilizing surface mutation identified using FoldX (A76D) and a mutation identified using PROSS (T197Y), produced D-SerFS (LSQED-Y), a variant of the sensor that exhibited high thermostability (Tm = 79.0 ± 0.3 °C) and an improved binding affinity for D-serine (*K*_D_ = 7.0 ± 0.4 μM), while maintaining specificity for D-serine.

Several stabilizing mutations remote from the binding site provided unexpected improvements in binding affinity. In particular, T197Y improved the binding affinity for all ligands while maintaining D-serine specificity. Such effects may be attributed to shifts in the conformational equilibrium of the binding protein. As the binding affinity of a protein for a given ligand may be considered in terms of the equilibria between the open or closed, and apo or bound states, manipulating these equilibria at positions distant from the binding site can lead to changes in affinity. Indeed, investigation of the T197Y mutation in the closed crystal structure of DalS (PDB 4DZ1) and a representative open conformation obtained from MD simulations, suggested that the mutation would stabilize the ligand bound (closed) conformation. The effect of the T197Y mutation on the relative populations of the open and closed conformations is also consistent with the observed decrease in dynamic range (13% in LSQED *vs*. 7% in LSQED-Y). This suggests a possible trade-off between improved binding affinity (*via* stabilization of the closed conformation) and dynamic range in FRET sensor design. In this instance, the slightly reduced dynamic range caused by the T197Y mutation, was significantly outweighed by the improved thermostability and affinity for D-serine of D-serFS. Future design studies on SBPs or other dynamic binders may directly exploit conformational changes between open and closed states as we have to design highly sensitive probes.

The repair of an unintended C-terminal truncation in the Venus fluorescent protein of D-serFS produced a full-length variant of the sensor that displayed reasonable dynamic range *in-vitro* (DR = ~ 14%) while maintaining similar binding affinity (*K*_D_ for D-serine = 4.8 ± 0.2 μM) and thermostability (Tm = 78.8 ± 1.0 °C) as the truncated variant. Testing the full-length variant of D-serFS in acute hippocampal rat brain slices demonstrated that the sensor is responsive to D-serine and its concentration changes in the environment of acute slices and is compatible with 2PE fluorescence microscopy. Assuming that the reduction in the response of D-serFS to saturating concentrations of D-serine in acute brain slices (Fig. **6D**) compared to responses in D-serine free solution (Fig. **6B**) is due to the extracellular ambient D-serine in the brain tissue and its binding to D-serFS, an estimate of the resting D-serine concentration can be obtained using the two responses and the *K*_D_ (see Zhang et al., 2018, Whitfield et al., 2015). Doing so we can estimate the resting concentration of D-serine to be ~ 5-6 μM. This is in line with previous reports using microdialysis, which also measured extracellular D-serine concentrations in the low micromolar range *in vivo* (rat striatum ~ 8 μM (Ciriacks and Bowser, 2004), rat frontal cortex ~ 6 μM (Matsui et al., 1995), mouse barrel cortex 4.2 μM (Takata et al., 2011). Compared to the reported D-serine resting levels, the apparent *K*_D_ of D-serFS (6.7 ± 0.5 μM depending on method) is ideal to detect both increases and decreases of the ambient D-serine concentrations in physiological settings and disease models. This was demonstrated by the sensitivity of D-serFS to the presence of the D-serine catabolizing enzyme DAAO in brain slices.

### Significance

In this work, we have used computational protein design to engineer an existing glycine/D - alanine binding protein (DalS) towards D-serine specificity and greater stability for use in a FRET-based biosensor for D-serine (full-length D-serFS). We demonstrated that D-serFS can be used to detect changes of extracellular D-serine levels in rat brain tissue, providing a new tool for the *in-situ* and potentially *in-vivo* study of the transmitter. This work highlights the utility of computational design tools in engineering naturally occurring binding proteins towards novel specificities, particularly where low-throughput experimental tests of affinity are required to discern the subtle effects of binding site mutations on specificity.

## Materials and Methods

### DNA cloning and mutagenesis

The DalS (Osborne et al., 2012) wild-type gene was synthesized (GeneArt) for sensor cloning, codon-optimized for expression in *Escherichia coli*. Sensor constructs were cloned into a vector backbone denoted as ‘pDOTS10’. This utilizes a vector system described previously (Okumoto et al., 2005) (Addgene Plasmid #13537), which contains a pRSET backbone with an N-terminal 6xHis tag and the insertion of a biotin tag from the PinPoint™ Xa-1 Vector (Promega, USA) in between the His tag and the first fluorescent protein. Endogenous *SapI* sites were removed from the ECFP-Venus cassette, and the binding protein gene YbeJ was replaced with a *SapI* linker (ATCAgaagagcactgcatggtGCGGCCGCcaccactctcgctcttcCCTC) designed around the method of Golden Gate cloning (Engler et al., 2008), whereby *SapI* restriction sites (GCTCTTC/GAAGAGC) are used to clone in a gene of interest. Reciprocal *SapI* sites were added to the 5’ and 3’ ends of the DalS gene by PCR for subsequent cloning into pDOTS10. Mutations were introduced using a combination of long mutagenesis primers and T7 promoter/terminator primers to create gene fragments with >40 base pair overlaps assembled *via* Gibson Assembly (Gibson et al., 2009). These variants were cloned into a new vector backbone (pETMCSIII), retaining the fluorescent proteins and biotin tag. Cloning to repair the truncation of the Venus fluorescent protein in D-serFS was performed using amplification primers to generate three gene fragments corresponding to 1) the N-terminal 6xHis tag, biotin purification tag, and ECFP, 2) the D-serFS binding protein, and 3) full-length Venus and the pDOTS10 vector backbone. The fragments generated had >40 bp overlaps and were assembled *via* Gibson Assembly (Gibson et al., 2009).

### Protein expression and purification

All proteins were expressed through transformation into BL21(DE3) *E. coli* cells and grown for 72 – 96 h at 18 °C in 1 L autoinducing medium (yeast extract, 5 g; tryptone, 20 g; NaCl, 5 g; KH_2_PO_4_, 3 g; Na_2_HPO_4_, 6 g in 1 L of water to which 10 ml autoclaved 60% (v/v) glycerol, 5 ml autoclaved 10% (w/v) glucose, 25 ml autoclaved 8% (w/v) lactose were added) supplemented with 100 mg ampicillin. The full expression of the fluorescent protein constructs needed to be monitored by observing the ECFP/Venus spectra over time, with Venus emission typically peaking at 72 h of expression at 18 °C.

Cells were harvested through centrifugation and the pellet was stored at −20 °C prior to purification. For purification, the pellet (frozen or otherwise) was suspended in buffer A (50 mM phosphate, 300 mM NaCl, 20 mM imidazole, pH 7.5), lysed by sonication, re-centrifuged at high speed (13,500 r.p.m. for 45 min at 4 °C) and the clarified supernatant was collected. The supernatant was loaded onto a 5 mL Ni-NTA/His-trap column pre-equilibrated in buffer A, washed with 10 column volumes of buffer A, 5 column volumes of 10% buffer B, and eluted with 100% buffer B (50 mM phosphate, 300 mM NaCl, 250 mM imidazole, pH 7.5). The eluted protein was dialyzed against 3 exchanges of 4 L of buffer C (20 mM phosphate, 200 mM NaCl, pH 7.5) at 4 °C. The dialyzed protein was further purified using a HiLoad 26/600 Superdex 200 pg SEC column using buffer C.

### Fluorescence assays

Fluorescence titrations were performed on a Varian Cary Eclipse using a Quartz narrow volume fluorescence cuvette. Samples underwent excitation at 433 nm and were scanned over a range of 450 nm – 570 nm for full spectra analysis. ECFP/Venus ratios were determined using peak wavelength values of 475 nm (ECFP) and 530 nm (Venus). Temperature dependent measurements were obtained using an Applied Photophysics Chirascan™ fluorescence photomultiplier with 433/3 nm excitation and peak fluorescence measured at 475 nm and 530 nm. *K*_D_ values were determined by fitting curves with the following equation:

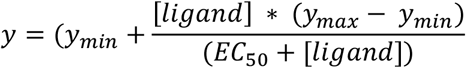

### Computational assessment of mutations

Mutations were assessed by first creating the mutation using the BuildModel command in FoldX (Schymkowitz et al., 2005), using the DalS crystal structure (Osborne et al., 2012) (PDB 4DZ1) that had undergone the FoldX Repair process and had the bound ligand removed. Prior to Schrödinger Glide (Friesner et al., 2004) docking, the Protein Preparation Wizard (Madhavi Sastry et al., 2013) was used to assign bond orders, generate protonation states and optimize the hydrogen bonding network at pH 7.4. Minimization of the complex using the OPLS3e (Roos et al., 2019) force field was completed with heavy atoms restrained to 0.30 Å of the input structure. Ligands were downloaded in SDF format and LigPrep was used to generate possible ionization states of the ligand at pH 7.4 ± 2.0. In Receptor Grid Generation, the default scaling factor of the Van der Waals radii of receptor atoms (1.0) and partial charge cut-off (0.25) were used. Rotatable receptor hydroxyl and thiol groups were allowed to rotate. The default scaling factor of the Van der Waals radii for the ligand (0.80) and partial charge cut-off (0.15) were used. In Glide, the standard precision (SP) docking method was used with ligand sampling set to the flexible setting, the option of adding Epik state penalties to the docking score included, and post-docking minimization performed. For Prime MM-GBSA (Rapp et al., 2011) calculations, the VSGB solvent model and OPLS3e force field were used in the computation of binding free energy. No residues were allowed to flex during minimization. For the identification of stabilising mutations using FoldX (Schymkowitz et al., 2005), the PositionScan command was used to mutate the targeted positions to each of the other amino acids, using the DalS crystal structure that had undergone the FoldX Repair process and had the bound ligand removed as the input.

### Molecular dynamics simulations and analysis

The crystal structure of DalS (Osborne et al., 2012) (PDB 4DZ1) was prepared for MD simulation by first building in missing residues using the mutagenesis wizard in PyMOL and removing the D-alanine ligand. The repaired structure was then capped with an N-terminal acetyl group and C-terminal N-methyl amide group. The protein was solvated in a rhombic dodecahedral box with 13 634 TIP3P water molecules, 37 sodium ions and 41 chloride ions (200 mM salt solution). Simulations were performed using GROMACS version 2018.3 (Abraham et al., 2015) using the CHARMM36m forcefield (Best et al., 2012). Long-range electrostatics were treated with the Particle Mesh Ewald method and the Van der Waals cut-off was set to 1.2 nm. The temperature was coupled to a virtual water bath at 300 K with a Bussi-Donadio-Parrinello thermostat. The Berendsen barostat was used during equilibrations with a time constant of 2 fs. Production runs were pressure coupled with a Parrinello-Rahman barostat with a time constant of 10 fs. A 2 fs time step was used throughout. Simulations were initially equilibrated with a 2 ns (1 000 000 steps) simulation and production runs were performed for 100 ns each (50 000 000 steps). 10 replicates of the equilibration and production simulations were performed. Cα–Cα distances and radius of gyration for each frame (sampled every 0.1 ns) was calculated using the ProDy package (Bakan et al., 2011). All replicates were concatenated prior to clustering analysis. Clustering was performed using the gromos method for clustering with an RMSD cut-off of 0.2 nm. The middle structure for the largest cluster was taken as the representative open conformation.

### Two-photon excitation (2PE) sensor imaging

Experiments were performed as previously described (Zhang et al., 2018, Whitfield et al., 2015) with minor changes. We used a FV10MP imaging system (Olympus) optically linked to a femtosecond pulse laser (Vision S, Coherent) equipped with a 25x (NA 1.05) objective (Olympus). For titrations in solution, D-serFS was imaged in a meniscus of PBS at a laser power of 3 mW and increasing amounts of D-serFS (in PBS) were added. For slice experiments (see below for further details), the laser power was adjusted for depth in the tissue to obtain, on average, a fluorescence intensity corresponding to that of 2-3 mW laser power at the slice surface. The excitation wavelength was 800 nm throughout. Fluorescence of ECFP and Venus fluorescent protein was separated using appropriate band pass filters and dichroic mirrors and detected with photomultipliers connected to a single photon counting board (Picoharp, Picoquant). Their arrival times were recorded using Symphotime 1.5 software (Picoquant). Offline analysis was performed using OriginPro 2017 (OriginLab) and custom written scripts in Matlab R2017a (Mathworks). The ratio of ECFP and Venus fluorescence intensity (R) was calculated from the number of photons detected by the respective detectors in time bins of ~ 800 ms. The photon count rate 11.5 ns to 12.5 ns after the laser pulse (81-82 MHz repetition rate) was subtracted to reduce the contribution of photons not originating from D-serFS to analysis.

### Brain slice preparation

Acute hippocampal slices were prepared from three- to five-week-old male Wistar rats as previously described (Anders et al., 2014, Zhang et al., 2018). All animals used in this study were housed under 12 h light/dark conditions and were allowed ad libitum access to food and water. Briefly, 350 μm thick horizontal slices containing hippocampal formation were obtained in full compliance with national and institutional regulations (Landesamt für Natur, Umwelt und Verbraucherschutz Nordrhein-Westfalen and University of Bonn Medical School) and guidelines of the European Union on animal experimentation. Slices were prepared in an ice-cold slicing solution containing (in mM): NaCl 60, sucrose 105, KCl 2.5, MgCl_2_ 7, NaH_2_PO_4_ 1.25, ascorbic acid 1.3, sodium pyruvate 3, NaHCO_3_ 26, CaCl_2_ 0.5, and glucose 10 (osmolarity 305–310 mOsm) and kept in the slicing solution at 34 °C for 15 min before being stored at room temperature in an extracellular solution containing (in mM) NaCl 131, KCl 2.5, MgSO_4_ 1.3, NaH_2_PO_4_ 1.25, NaHCO_3_ 21, CaCl_2_ 2, and glucose 10 (osmolarity 297–303 mOsm, pH adjusted to 7.4). This solution was also used for recordings. Slices were allowed to rest for at least 50 min. All solutions were continuously bubbled with 95% O_2_/ 5% CO_2_.

### Full-length D-serFS imaging in acute slices

D-serFS was handled and immobilized in acute hippocampal slices as described before (Zhang et al., 2018, Whitfield et al., 2015). For long-term storage and transport full-length D-serFS was lyophilized and stored for up to 2 months at ambient temperature (for shipping) and 4° C (storage until use). Before experiments, the sensor was reconstituted in water, agitated for 30 min at room temperature before the buffer was changed to PBS (pH 7.4) with a PD-10 desalting column (GE Healthcare). Full-length D-serFS was then concentrated to 200-300 μM using centricons (Vivaspin 500, 10 kDa cutoff, Sartorius Stedim Biotech). For anchoring of optical sensors in brain tissue, cell surfaces within acute slices were biotinylated using a previously published procedure (Zhang et al., 2018, Whitfield et al., 2015). Briefly, the slice storage solution was supplemented with 50 μM Sulfo-NHS EZ Link Biotin (Thermo Fisher) for 45 min before washing and further storage. Slices were transferred to a submersion-type recording chamber and superfused with extracellular solution at 34 °C. For injections of full-length D-serFS into the tissue, patch clamp pipettes (2-4 MΩ when filled with PBS saline) were backfilled with PBS (pH 7.4) to which 100 μM full-length D-serFS and 12.5 μM streptavidin (Life Technologies) had been added. The pipette was inserted ~70 μm deep into the tissue under visual control and full-length D-serFS was pressure-injected. Full-length D-serFS-injected acute slices were allowed to recover for 15 minutes before recordings. To reduce the potential effect of scattering of ECFP and Venus fluorescence, we have performed all imaging experiments at a depth of 50-70 μm below the slice surface. Exogenous D-serine was applied via the bath perfusion (1 mM). The following inhibitors were present throughout these experiments to block excitatory synaptic transmission and action potential firing: TTX (1 μM, Sigma Aldrich), D-APV (50 μM, Abcam) and NBQX (10 μM, Abcam). In a subset of experiments, slices were incubated for at least 45 min with DAAO (~ 0.2 U/ml, Sigma-Aldrich) at room temperature for D-serine degradation (Papouin et al., 2012). In this case, DAAO (~ 0.2 U/ml) was also added in the recording solution.

### Statistics

Data are reported as mean ± s.e.m. where *n* is the number of independently performed experiments, unless stated otherwise. For *in-vitro* experiments, independent experiments are defined as technical repeats performed using the same batch of purified sensor.

## Acknowledgements

V.V., and J.A.M. acknowledge financial support from an Australian Government Research Training Program Scholarship. Research was funded by an ARC Discovery Project awarded to C.J.J. Research in the Fleishman lab was funded by the European Research Council (815379), the Israel Science Foundation (1844), the Milner Foundation and a charitable donation from Sam Switzer and family. The work was further supported by the Human Frontiers Science Program (HFSP; RGY0084/2012 to C.H., H.J. and C.J.J.), the German Academic Exchange Service (DAAD-Go8) Travel Fellowship (to C.H. and C.J.J.), the NRW-Rückkehrerprogramm (C.H.) and German Research Foundation (DFG; SFB1089 B03, SPP1757 HE6949/1, FOR2795 and HE6949/3 to C.H.; SPP1757 young investigator grant to P.U.).

## Contributions

J.H.W., V.V., J.A.M., O.K., S.J.F., and C.J.J. designed, produced and analyzed the sensor. P.U., B.B., L.K., H.M., H.H. and C.H. performed and analyzed all experiments using two-photon excitation. H.J., C.J.J. and C.H. conceptualized the study. C.J.J. and V.V. wrote the initial manuscript, which all authors subsequently reviewed and edited.

## References

Abraham, M. J., Murtola, T., Schulz, R., Páll, S., Smith, J. C., Hess, B. & Lindahl, E. 2015. GROMACS: High performance molecular simulations through multi-level parallelism from laptops to supercomputers. SoftwareX, 1-2, 19–25.

Anders, S., Minge, D., Griemsmann, S., Herde, M. K., Steinhäuser, C. & Henneberger, C. 2014. Spatial properties of astrocyte gap junction coupling in the rat hippocampus. Philosophical Transactions of the Royal Society B: Biological Sciences, 369, 20130600.

Bajar, B. T., Wang, E. S., Zhang, S., Lin, M. Z. & Chu, J. 2016. A Guide to Fluorescent Protein FRET Pairs. Sensors (Basel), 16.

Bakan, A., Meireles, L. M. & Bahar, I. 2011. ProDy: Protein Dynamics Inferred from Theory and Experiments. Bioinformatics, 27, 1575–1577.

Basu, A. C., Tsai, G. E., Ma, C. L., Ehmsen, J. T., Mustafa, A. K., Han, L., Jiang, Z. I., Benneyworth, M. A., Froimowitz, M. P., Lange, N., Snyder, S. H., Bergeron, R. & Coyle, J. T. 2009. Targeted disruption of serine racemase affects glutamatergic neurotransmission and behavior. Molecular psychiatry, 14, 719–727.

Best, R. B., Zhu, X., Shim, J., Lopes, P. E. M., Mittal, J., Feig, M. & Mackerell, A. D. 2012. Optimization of the Additive CHARMM All-Atom Protein Force Field Targeting Improved Sampling of the Backbone ϕ, Ψ and Side-Chain χ1 and χ2 Dihedral Angles. Journal of Chemical Theory and Computation, 8, 3257–3273.

Beyene, A. G., Yang, S. J. & Landry, M. P. 2019. Review Article: Tools and trends for probing brain neurochemistry. Journal of vacuum science & technology. A, Vacuum, surfaces, and films : an official journal of the American Vacuum Society, 37, 040802–040802.

Bliss, T. V. & Collingridge, G. L. 1993. A synaptic model of memory: long-term potentiation in the hippocampus. Nature, 361, 31–9.

Bliss, T. V. P. & Cooke, S. F. 2011. Long-term potentiation and long-term depression: a clinical perspective. Clinics (Sao Paulo, Brazil), 66 Suppl 1,3–17.

Bus, O., Rudat, J. & Ochsenreither, K. 2018. FoldX as Protein Engineering Tool: Better Than Random Based Approaches? Computational and Structural Biotechnology Journal, 16, 25–33.

Ciriacks, C. M. & Bowser, M. T. 2004. Monitoring d-Serine Dynamics in the Rat Brain Using Online Microdialysis-Capillary Electrophoresis. Analytical Chemistry, 76, 6582–6587.

Clifton, B. E., Kaczmarski, J. A., Carr, P. D., Gerth, M. L., Tokuriki, N. & Jackson, C. J. 2018. Evolution of cyclohexadienyl dehydratase from an ancestral solute-binding protein. Nature Chemical Biology, 14, 542–547.

Dai, X., Zhou, E., Yang, W., Zhang, X., Zhang, W. & Rao, Y. 2019. D-Serine made by serine racemase in Drosophila intestine plays a physiological role in sleep. Nature Communications, 10, 1986.

Dale, N., Hatz, S., Tian, F. & Llaudet, E. 2005. Listening to the brain: microelectrode biosensors for neurochemicals. Trends in Biotechnology, 23, 420–428.

Engler, C., Kandzia, R. & Marillonnet, S. 2008. A One Pot, One Step, Precision Cloning Method with High Throughput Capability. PLOS ONE, 3, e3647.

Friesner, R. A., Banks, J. L., Murphy, R. B., Halgren, T. A., Klicic, J. J., Mainz, D. T., Repasky, M. P., Knoll, E. H., Shelley, M., Perry, J. K., Shaw, D. E., Francis, P. & Shenkin, P. S. 2004. Glide: A New Approach for Rapid, Accurate Docking and Scoring. 1. Method and Assessment of Docking Accuracy. Journal of Medicinal Chemistry, 47, 1739–1749.

Furukawa, H. & Gouaux, E. 2003. Mechanisms of activation, inhibition and specificity: crystal structures of the NMDA receptor NR1 ligand-binding core. The EMBO Journal, 22, 2873–2885.

Ganesana, M., Lee, S. T., Wang, Y. & Venton, B. J. 2017. Analytical Techniques in Neuroscience: Recent Advances in Imaging, Separation, and Electrochemical Methods. Analytical chemistry, 89, 314–341.

Gibson, D. G., Young, L., Chuang, R.-Y., Venter, J. C., Hutchison, C. A. & Smith, H. O. 2009. Enzymatic assembly of DNA molecules up to several hundred kilobases. Nature Methods, 6, 343–345.

Goldenzweig, A., Goldsmith, M., Hill, S. E., Gertman, O., Laurino, P., Ashani, Y., Dym, O., Unger, T., Albeck, S., Prilusky, J., Lieberman, R. L., Aharoni, A., Silman, I., Sussman, J. L., Tawfik, D. S. & Fleishman, S. J. 2016. Automated Structure- and Sequence-Based Design of Proteins for High Bacterial Expression and Stability. Mol Cell, 63, 337–346.

Hardingham, G. E. & Bading, H. 2010. Synaptic versus extrasynaptic NMDA receptor signalling: implications for neurodegenerative disorders. Nature Reviews Neuroscience, 11, 682–696.

Hashimoto, A., Nishikawa, T., Hayashi, T., Fujii, N., Harada, K., Oka, T. & Takahashi, K. 1992. The presence of free D-serine in rat brain. FEBS Lett, 296, 33–6.

Henneberger, C., Papouin, T., Oliet, S. H. R. & Rusakov, D. A. 2010. Long-term potentiation depends on release of d-serine from astrocytes. Nature, 463, 232–236.

Kaczmarski, J. A., Mahawaththa, M. C., Feintuch, A., Clifton, B. E., Adams, L. A., Goldfarb, D., Otting, G. & Jackson, C. J. 2020. Altered conformational sampling along an evolutionary trajectory changes the catalytic activity of an enzyme. Nature Communications, 11, 5945.

Kaczmarski, J. A., Mitchell, J. A., Spence, M. A., Vongsouthi, V. & Jackson, C. J. 2019. Structural and evolutionary approaches to the design and optimization of fluorescence-based small molecule biosensors. Current Opinion in Structural Biology, 57, 31–38.

Labrie, V., Fukumura, R., Rastogi, A., Fick, L. J., Wang, W., Boutros, P. C., Kennedy, J. L., Semeralul, M. O., Lee, F. H., Baker, G. B., Belsham, D. D., Barger, S. W., Gondo, Y., Wong, A. H. & Roder, J. C. 2009. Serine racemase is associated with schizophrenia susceptibility in humans and in a mouse model. Hum Mol Genet, 18, 3227–43.

Liu, S., Liu, Q., Tabuchi, M. & Wu, M. N. 2016. Sleep Drive Is Encoded by Neural Plastic Changes in a Dedicated Circuit. Cell, 165, 1347–1360.

Lyne, P. D., Lamb, M. L. & Saeh, J. C. 2006. Accurate Prediction of the Relative Potencies of Members of a Series of Kinase Inhibitors Using Molecular Docking and MM-GBSA Scoring. Journal of Medicinal Chemistry, 49, 4805–4808.

Madeira, C., Lourenco, M. V., Vargas-Lopes, C., Suemoto, C. K., BrandãO, C. O., Reis, T., Leite, R. E. P., Laks, J., Jacob-Filho, W., Pasqualucci, C. A., Grinberg, L. T., Ferreira, S. T. & Panizzutti, R. 2015. d-serine levels in Alzheimer’s disease: implications for novel biomarker development. Translational Psychiatry, 5, e561–e561.

Madhavi Sastry, G., Adzhigirey, M., Day, T., Annabhimoju, R. & Sherman, W. 2013. Protein and ligand preparation: parameters, protocols, and influence on virtual screening enrichments. Journal of Computer-Aided Molecular Design, 27, 221–234.

Matsui, T.-A., Sekiguchi, M., Hashimoto, A., Tomita, U., Nishikawa, T. & Wada, K. 1995. Functional Comparison of d-Serine and Glycine in Rodents: The Effect on Cloned NMDA Receptors and the Extracellular Concentration. Journal of Neurochemistry, 65, 454–458.

Mohd Zain, Z., Ab Ghani, S. & O’Neill, R. D. 2012. Amperometric microbiosensor as an alternative tool for investigation of D-serine in brain. Amino Acids, 43, 1887–94.

Mothet, J.-P., Parent, A. T., Wolosker, H., Brady, R. O., Linden, D. J., Ferris, C. D., Rogawski, M. A. & Snyder, S. H. 2000. D-Serine is an endogenous ligand for the glycine site of the *N*-methyl-D-aspartate receptor. Proceedings of the National Academy of Sciences, 97, 4926–4931.

Okumoto, S., Looger, L. L., Micheva, K. D., Reimer, R. J., Smith, S. J. & Frommer, W. B. 2005. Detection of glutamate release from neurons by genetically encoded surface-displayed FRET nanosensors. Proceedings of the National Academy of Sciences of the United States of America, 102, 8740.

Osborne, S. E., Tuinema, B. R., Mok, M. C., Lau, P. S., Bui, N. K., Tomljenovic-Berube, A. M., Vollmer, W., Zhang, K., Junop, M. & Coombes, B. K. 2012. Characterization of DalS, an ATP-binding cassette transporter for D-alanine, and its role in pathogenesis in Salmonella enterica. J Biol Chem, 287, 15242–50.

Panatier, A., Theodosis, D. T., Mothet, J.-P., Touquet, B., Pollegioni, L., Poulain, D. A. & Oliet, S. H. R. 2006. Glia-Derived d-Serine Controls NMDA Receptor Activity and Synaptic Memory. Cell, 125, 775–784.

Papouin, T., Dunphy, J. M., Tolman, M., Dineley, K. T. & Haydon, P. G. 2017a. Septal Cholinergic Neuromodulation Tunes the Astrocyte-Dependent Gating of Hippocampal NMDA Receptors to Wakefulness. Neuron, 94, 840–854.e7.

Papouin, T., Henneberger, C., Rusakov, D. A. & Oliet, S. H. R. 2017b. Astroglial versus Neuronal D-Serine: Fact Checking. Trends in Neurosciences, 40, 517–520.

Papouin, T., Ladepeche, L., Ruel, J., Sacchi, S., Labasque, M., Hanini, M., Groc, L., Pollegioni, L., Mothet, J. P. & Oliet, S. H. 2012. Synaptic and extrasynaptic NMDA receptors are gated by different endogenous coagonists. Cell, 150, 633–46.

Pernot, P., Mothet, J.-P., Schuvailo, O., Soldatkin, A., Pollegioni, L., Pilone, M., Adeline, M.-T., Cespuglio, R. & Marinesco, S. 2008. Characterization of a Yeast d-Amino Acid Oxidase Microbiosensor for d-Serine Detection in the Central Nervous System. Analytical Chemistry, 80, 1589–1597.

Pollegioni, L. & Sacchi, S. 2010. Metabolism of the neuromodulator D-serine. Cell Mol Life Sci, 67, 2387–404.

Popiolek, M., Tierney, B., Steyn, S. J. & Devivo, M. 2018. Lack of Effect of Sodium Benzoate at Reported Clinical Therapeutic Concentration on d-Alanine Metabolism in Dogs. ACS Chemical Neuroscience, 9, 2832–2837.

Rapp, C., Kalyanaraman, C., Schiffmiller, A., Schoenbrun, E. L. & Jacobson, M. P. 2011. A molecular mechanics approach to modeling protein-ligand interactions: relative binding affinities in congeneric series. Journal of chemical information and modeling, 51, 2082–2089.

Roos, K., Wu, C., Damm, W., Reboul, M., Stevenson, J. M., Lu, C., Dahlgren, M. K., Mondal, S., Chen, W., Wang, L., Abel, R., Friesner, R. A. & Harder, E. D. 2019. OPLS3e: Extending Force Field Coverage for Drug-Like Small Molecules. Journal of Chemical Theory and Computation, 15, 1863–1874.

Schell, M. J., Molliver, M. E. & Snyder, S. H. 1995. D-serine, an endogenous synaptic modulator: localization to astrocytes and glutamate-stimulated release. Proceedings of the National Academy of Sciences, 92, 3948–3952.

Schymkowitz, J., Borg, J., Stricher, F., Nys, R., Rousseau, F. & Serrano, L. 2005. The FoldX web server: an online force field. Nucleic acids research, 33, W382–W388.

Shippenberg, T. S. & Thompson, A. C. 2001. Overview of microdialysis. Current protocols in neuroscience, Chapter 7, Unit7.1-Unit7.1.

Takata, N., Mishima, T., Hisatsune, C., Nagai, T., Ebisui, E., Mikoshiba, K. & Hirase, H. 2011. Astrocyte Calcium Signaling Transforms Cholinergic Modulation to Cortical Plasticity In Vivo. The Journal of Neuroscience, 31, 18155–18165.

Tokuriki, N., Stricher, F., Schymkowitz, J., Serrano, L. & Tawfik, D. S. 2007. The Stability Effects of Protein Mutations Appear to be Universally Distributed. Journal of Molecular Biology, 369, 1318–1332.

Tomita, J., Ueno, T., Mitsuyoshi, M., Kume, S. & Kume, K. 2015. The NMDA Receptor Promotes Sleep in the Fruit Fly, Drosophila melanogaster. PLOS ONE, 10, e0128101.

Whitfield, J. H., Zhang, W. H., Herde, M. K., Clifton, B. E., Radziejewski, J., Janovjak, H., Henneberger, C. & Jackson, C. J. 2015. Construction of a robust and sensitive arginine biosensor through ancestral protein reconstruction. Protein Science, 24, 1412–1422.

Wijma, H. J., Floor, R. J., Jekel, P. A., Baker, D., Marrink, S. J. & Janssen, D. B. 2014. Computationally designed libraries for rapid enzyme stabilization. Protein engineering, design & selection : PEDS, 27, 49–58.

Wiriyasermkul, P., Moriyama, S., Tanaka, Y., Kongpracha, P., Nakamae, N., Suzuki, M., Kimura, T., Mita, M., Sasabe, J. & Nagamori, S. 2020. D-Serine, an emerging biomarker of kidney diseases, is a hidden substrate of sodium-coupled monocarboxylate transporters. bioRxiv,2020.08.10.244822.

Wolosker, H., Balu, D. T. & Coyle, J. T. 2016. The Rise and Fall of the d-Serine-Mediated Gliotransmission Hypothesis. Trends in Neurosciences, 39, 712–721.

Wolosker, H., Blackshaw, S. & Snyder, S. H. 1999. Serine racemase: a glial enzyme synthesizing D-serine to regulate glutamate-N-methyl-D-aspartate neurotransmission. Proc Natl Acad Sci U S A, 96, 13409–14.

Yang, Y., Ge, W., Chen, Y., Zhang, Z., Shen, W., Wu, C., Poo, M. & Duan, S. 2003. Contribution of astrocytes to hippocampal long-term potentiation through release of D-serine. Proceedings of the National Academy of Sciences, 100, 15194.

Zhang, W. H., Herde, M. K., Mitchell, J. A., Whitfield, J. H., Wulff, A. B., Vongsouthi, V., Sanchez-Romero, I., Gulakova, P. E., Minge, D., Breithausen, B., Schoch, S., Janovjak, H., Jackson, C. J. & Henneberger, C. 2018. Monitoring hippocampal glycine with the computationally designed optical sensor GlyFS. Nature Chemical Biology, 14, 861–869.

