## Supporting_Information for "A computationally designed fluorescent biosensor for D-serine"

**Supplementary Table 1**. Scores computed by Glide (Friesner et al., 2004) and Prime MM-GBSA (Rapp et al., 2011). Glide scores (kcal/mol) and relative binding free energies (ΔG_bind_) predicted by Prime MM-GBSA for the top poses of D-alanine (D-ala) and D-serine (D-ser) are shown. The difference between the ΔG_bind_ of D-ser and D-ala represents the predicted specificity of each variant.

| **Variant** | **Glide Score (kcal/mol)** | |  | **ΔG_bind_ (kcal/mol)** | |  | **(D-ser – D-ala) ΔG_bind_ (kcal/mol)** |
| --- | --- | --- | --- | --- | --- | --- | --- |
|  | **D-ala** | **D-ser** |  | **D-ala** | **D-ser** |  |  |
| WT | -6.6 | -7.4 |  | -27.9 | -29.1 |  | -1.2 |
| LSQ | -6.0 | -7.9 |  | -22.5 | -27.8 |  | -5.3 |
| D216S | -5.5 | -5.0 |  | -15.1 | -21.0 |  | -5.9 |
| D216T | -6.0 | -5.9 |  | -13.3 | -14.8 |  | -1.5 |
| D216E | -6.0 | -6.3 |  | -12.0 | -20.8 |  | -8.8 |
| N115Q | -6.4 | -8.1 |  | -28.3 | -25.1 |  | 3.2 |
| N115S | -7.5 | -8.7 |  | -25.0 | -30.1 |  | -5.1 |
| N115T | -7.3 | -8.7 |  | -23.9 | -28.6 |  | -4.8 |
| Y173A | -6.1 | -7.5 |  | -20.1 | -23.7 |  | -3.6 |
| Y173F | -6.9 | -6.6 |  | -12.8 | -13.1 |  | -0.3 |

**Supplementary Table 2.** Change in free energy of folding (ΔΔG) calculated by both FoldX and Rosetta computational mutation scanning for all stabilising mutations predicted by FoldX PositionScan (grey) and PROSS (no shading). Mutations are ranked by increasing ΔΔG (kcal/mol) predicted by FoldX. Variants above the dashed line had a ΔΔG more negative than 2.5 FoldX standard deviations (< 1.15 kcal/mol) and were selected for experimental testing.

| **Variant** | **Predicted ΔΔG** |  |  | **ΔTm relative to LSQED (°C)** |
| --- | --- | --- | --- | --- |
|  | FoldX (kcal/mol) | Rosetta (R.e.u.) |  |  |
| A76D | -2.30 | -1.82 |  | 2.8^a^ |
| S208E | -1.94 | 8.22 |  | 1.2 |
| S60K^b^ | -1.60 | -0.26 |  | 1.3 |
| T172D | -1.41 | 4.78 |  | 6.0 |
| S60R | -1.31 | -0.33 |  | 14.0 |
| T197Y | -1.25 | -0.82 |  | 14.0 |
| N200K | -1.17 | -0.34 |  | 4.4 |
| H67G | -1.04 | -0.29 |  | - |
| A76S | -0.87 | -1.87 |  | - |
| G82A | -0.85 | -0.69 |  | - |
| Q242E | -0.85 | -0.40 |  | - |
| K199L | -0.80 | -0.60 |  | - |
| G246A | -0.78 | -0.62 |  | - |
| S61A | -0.67 | -1.12 |  | - |
| D150E | -0.65 | 2.51 |  | - |
| G246E | -0.65 | -0.50 |  | - |
| N32R | -0.63 | 1.07 |  | - |
| S245K | -0.50 | -1.62 |  | - |
| S245Q | -0.43 | -0.84 |  | - |
| A123K | -0.33 | -0.16 |  | - |
| Q88K | -0.20 | -0.49 |  | - |
| T73V | -0.09 | -0.78 |  | - |
| N136K | -0.09 | -0.18 |  | - |
| H125D | -0.04 | -0.59 |  | - |
| T176A | 0.04 | -0.41 |  | - |
| T176V | 0.04 | 2.52 |  | - |
| N133S | 0.09 | -1.16 |  | - |
| Q64K | 0.09 | -0.36 |  | - |
| E110D | 0.24 | -0.46 |  | - |
| S65E | 0.33 | -0.36 |  | - |
| N131D | 0.44 | 0.16 |  | - |
| I114T | 0.46 | -0.17 |  | - |
| L234D | 1.19 | -0.24 |  | - |
| L70C | 2.39 | 1.02 |  | - |

^a^ΔTm of A76D is relative to LSQE. All other experimentally determined ΔTm values are relative to LSQE/A76D (LSQED).

^b^S60K was predicted by both FoldX PositionScan and PROSS.

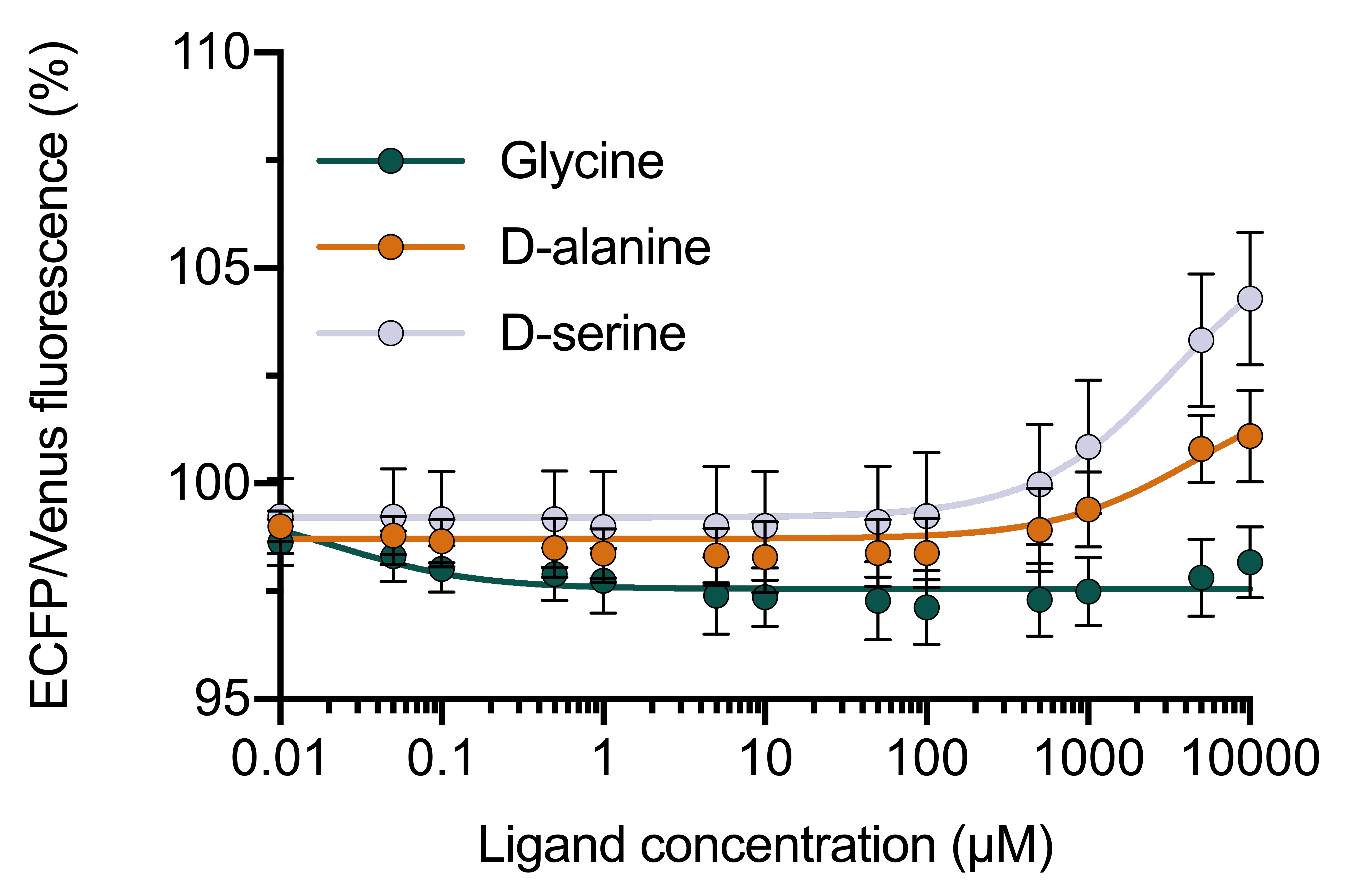

**Supplementary Figure 1.** Dose-response curves for F117**L**/A147**S**/Y148**V**/D216**E**/A76**D** (LSVED) with glycine, D-alanine and D-serine. Values are the (475 nm/530 nm) fluorescence ratio as a percentage of the same ratio for the apo sensor. No significant change is detected in response to glycine. The *K_D_* for D-alanine and D-serine are estimated to be > 4000 μM based on fitting curves with the following equation: $y=(y_{min}+\frac{\left[ ligand \right] * (y_{max}- y_{min})}{{(EC}_{50}+[ligand])}$ .

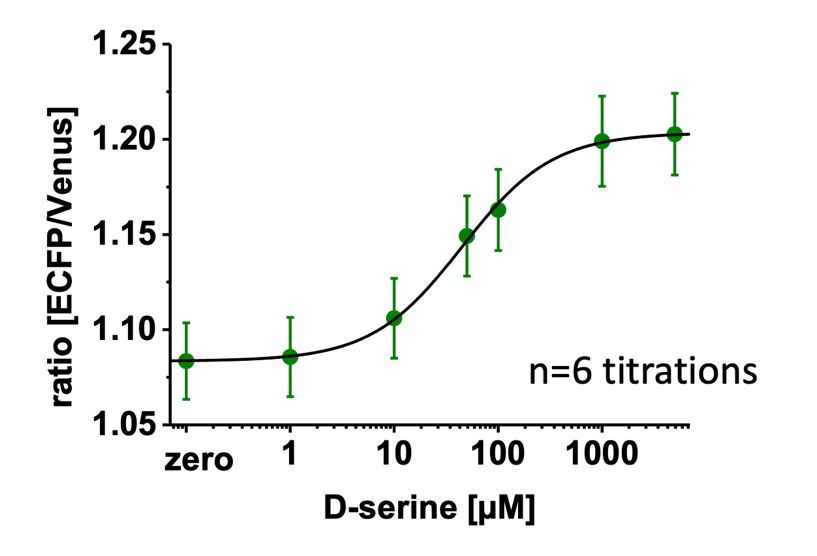

**Supplementary Figure 2.** Preliminary titration of the LSQ variant under two-photon excitation (2PE) fluorescence microscopy, demonstrating a reduced dynamic range (~ 10%) following lyophilisation and reconstitution compared to prior (~ 13%).

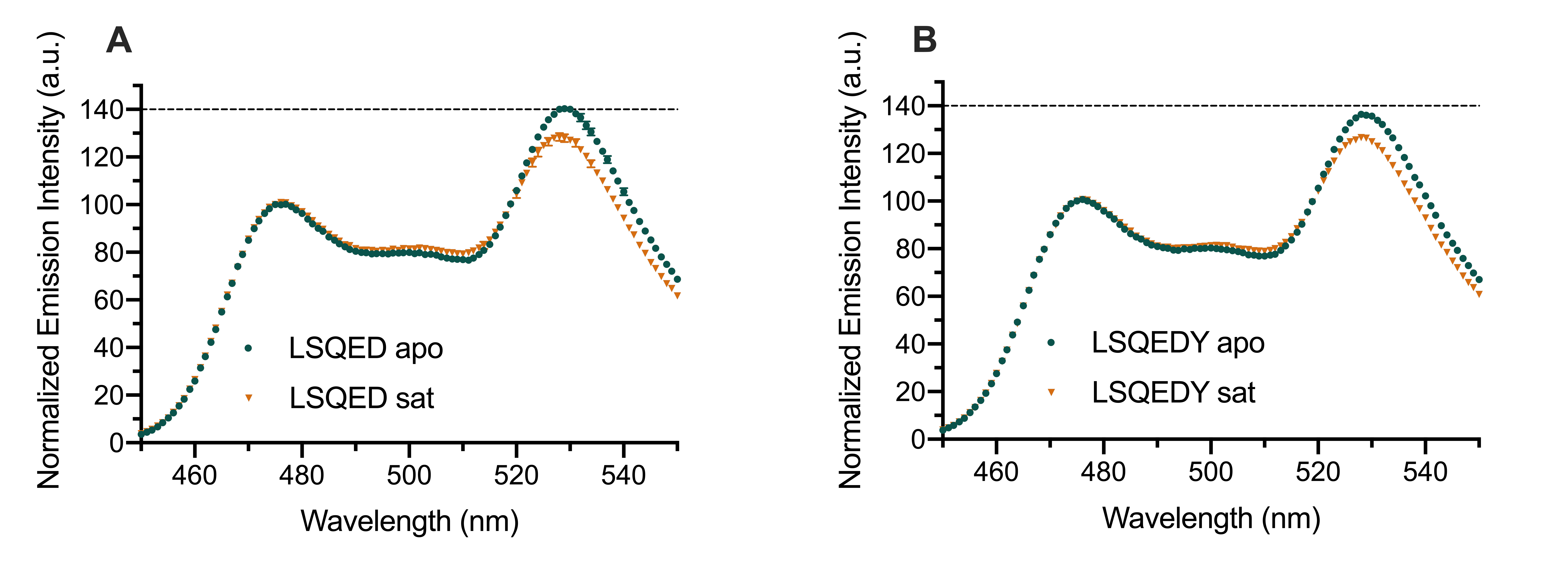

**Supplementary Figure 3.** Emission spectra (450 – 550 nm) of (A) LSQED and (B) LSQED-T197Y (LSQEDY) upon excitation of ECFP (λ_exc_ = 433 nm), normalised to the maximum emission intensity from ECFP (475 nm). For all sensor variants, the FRET efficiency decreases in response to saturation with D-serine (A, B; orange), leading to decreased emission from Venus (530 nm) relative to ECFP (475 nm). When comparing the apo states of LSQED and LSQEDY (A, B; dark green), it can be seen that the T197Y mutation results in a decreased Venus emission (lower FRET efficiency). This suggests a shift in the apo population of the sensor towards the spectral properties of the saturated, closed state and explains the decreased dynamic range of LSQEDY compared to LSQED. Values are mean ± s.e.m (n = 3).

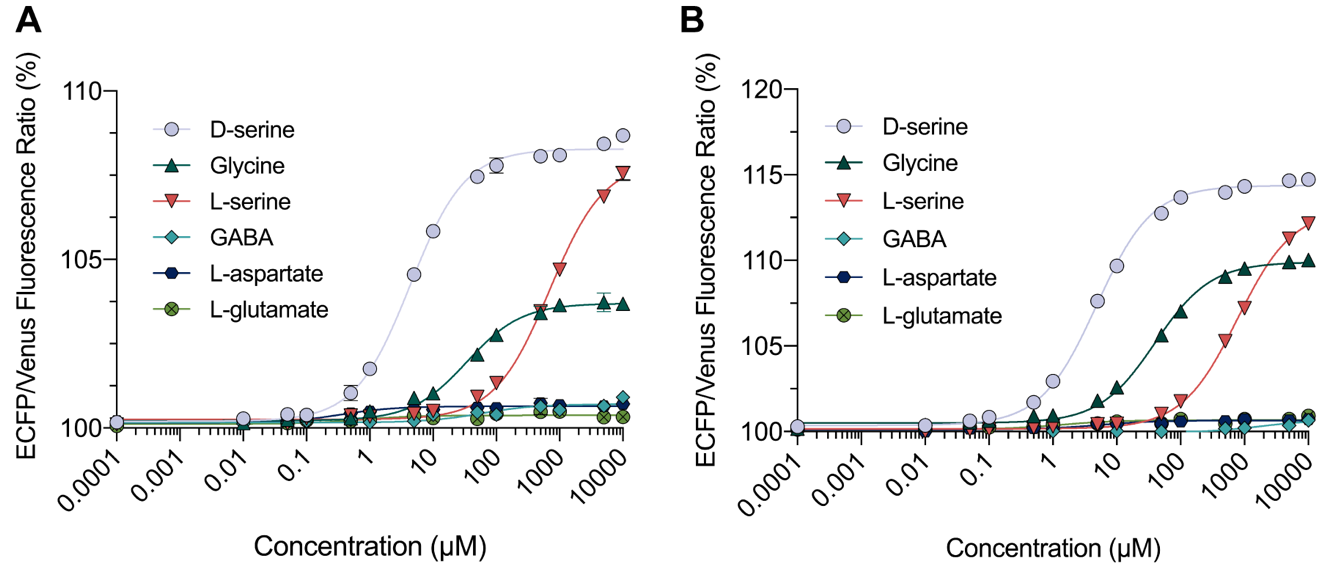

**Supplementary Figure 4.** Fluorescence titration of A) truncated D-serFS (LSQED+T197Y) and B) full-length D-serFS with D-serine, glycine, L-serine, GABA, L-aspartate, and L-glutamate.

No binding of GABA, L-aspartate or L-glutamate for either variant was detected. The purity of L-serine used was ~ 99%, thus, concentrations of 500 and 1000 μM of L-serine possibly contain D-serine at concentrations of ~ 5 and ~ 10 μM, respectively, causing an increase in the ECFP/Venus fluorescence ratio in the titration with L-serine for both sensor variants.

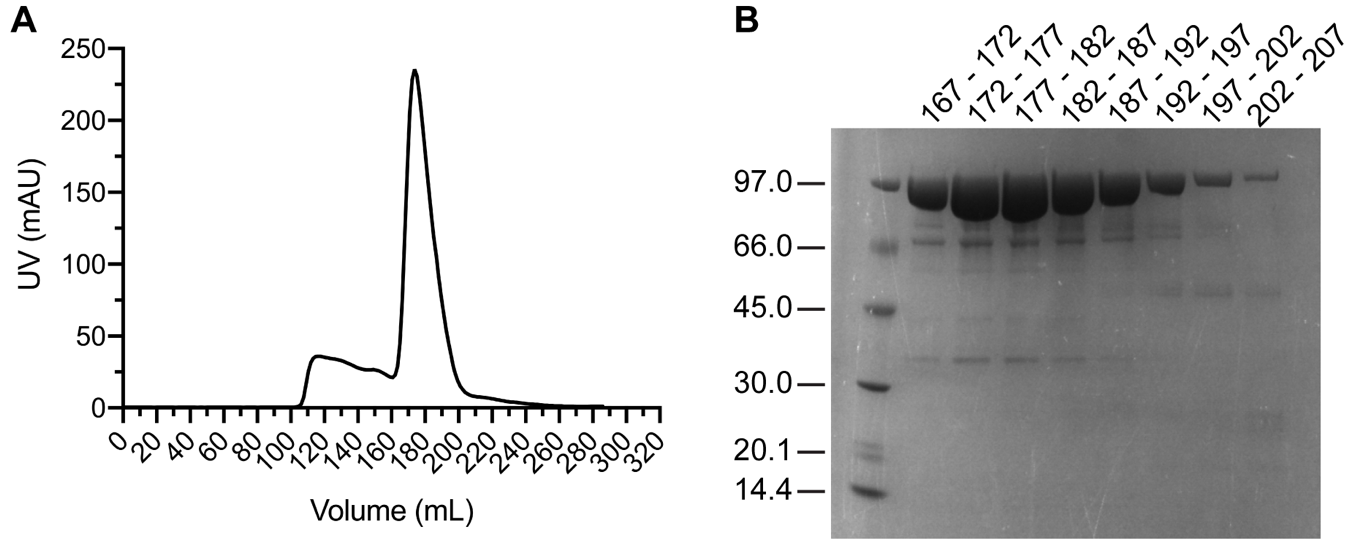

**Supplementary Figure 5** A) Chromatogram from size-exclusion chromatography (HiLoad 26/600 Superdex 200 pg) of D-SerFS following Ni^2+^-affinity chromatography. The main elution peak occurs between 167 – 207 mL. B) Eight 5 mL fractions from the elution peak (167 – 207 mL) were analysed for purity by SDS-PAGE. The most concentrated fractions occurred between 172 – 187 mL and contained minor impurities compared to the band corresponding to D-SerFS (97 kDa). These fractions were pooled for subsequent experiments.

**Supplementary Figure 6** Full amino-acid sequence of full-length D-Ser-FS

6xHis Tag, Biotin Purification Tag, ECFP, Linker 1, DalS LSQED-Y binding core, Linker 2, Venus

HHHHHHGMASMTGGQQMGRDLYDDDDKDPKLKVTVNGTAYDVDVDVDKSHENPMGTILFGGGTGGAPAPAAGGAGAGKAGEGEIPAPLAGTVSKILVKEGDTVKAGQTVLVLEAMKMETEINAPTDGKVEKVLVKERDAVQGGQGLIKIGDLELIEGSSGSDPGRMVSKGEELFTGVVPILVELDGDVNGHKFSVSGEGEGDATYGKLTLKFICTTGKLPVPWPTLVTTLTWGVQCFSRYPDHMKQHDFFKSAMPEGYVQERTIFFKDDGNYKTRAEVKFEGDTLVNRIELKGIDFKEDGNILGHKLEYNYISHNVYITADKQKNGIKANFKIRHNIEDGSVQLADHYQQNTPIGDGPVLLPDNHYLSTQSALSKDPNEKRDHMVLLEFVTAAGITLGMDELYKGGTGIMIVEGRTLNVAVSPASPPMLFKSADGKLQGIDLELFSSYCQSRHCKLNITEYDWDGMLGAVASGQADVAFSGISITDKRKKVIDFSEPYYINSLYLVSMANHKITLNNLNELNKYSIGYPRGMSQSDLIKNDLEPKGYYSLSKVKLYPTYNETMADLKNGNLDLAFIEEPVYFYFKNKKKMPIESRYVFKNVEQLGIAFKKGSPVRDDFNLWLKEQGPQKISGIVDSWMKLAGTGGMVSKGEELFTGVVPILVELDGDVNGHKFSVSGEGEGDATYGKLTLKLICTTGKLPVPWPTLVTTLGYGLQCFARYPDHMKQHDFFKSAMPEGYVQERTIFFKDDGNYKTRAEVKFEGDTLVNRIELKGIDFKEDGNILGHKLEYNYNSHNVYITADKQKNGIKANFKIRHNIEDGGVQLADHYQQNTPIGDGPVLLPDNHYLSYQSALSKDPNEKRDHMVLLEFVTAAGITLGMDELYK*

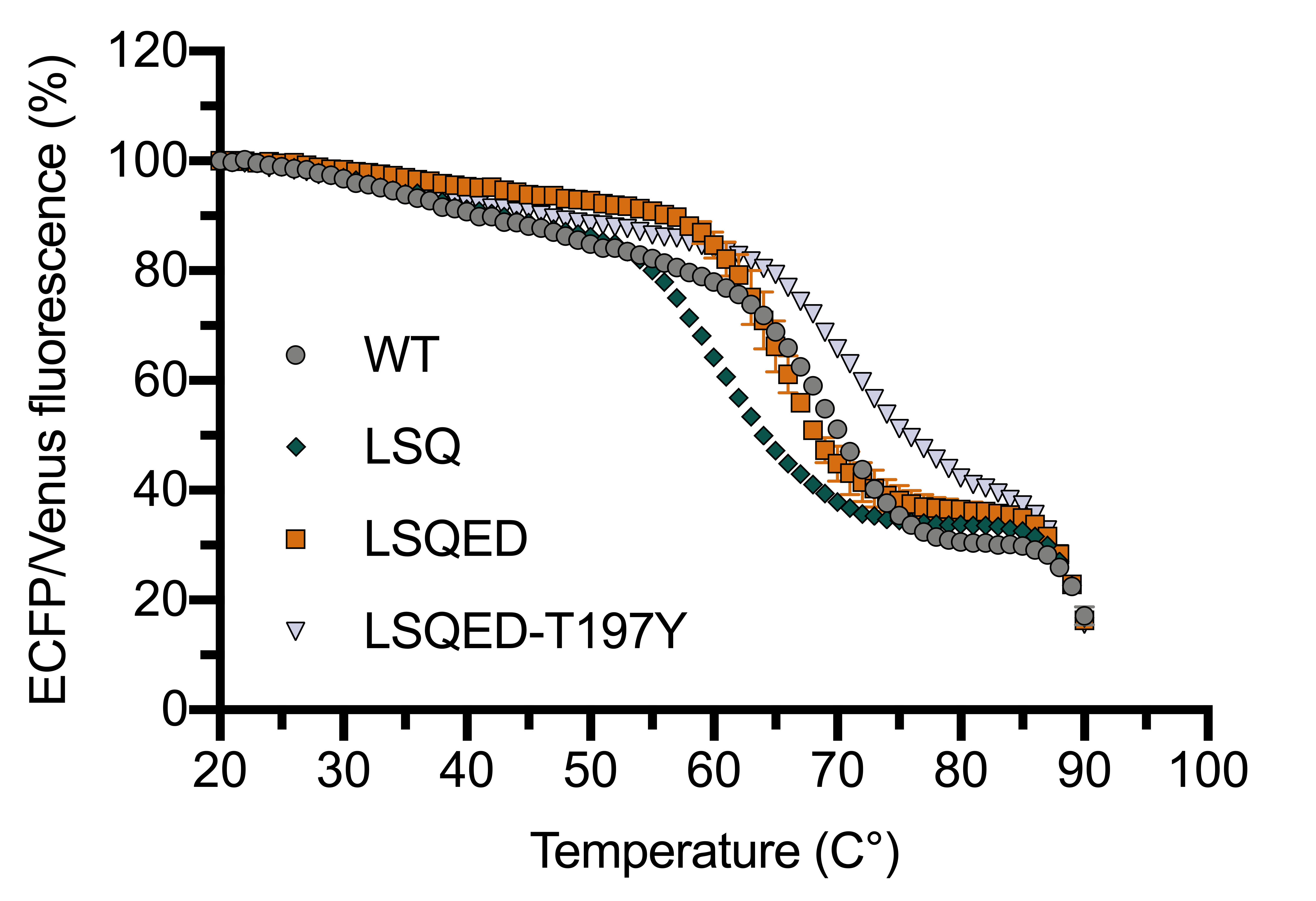

**Supplementary Figure 7**. The (475 nm/530 nm) fluorescence ratio as a function of increasing temperature (20 – 90 °C) for key variants in the engineering trajectory of D-serFS. Values are normalised as a percentage of the same ratio for the sensor at 20 °C and are represented as mean ± s.e.m. (n = 3). The first sigmoidal transition (corresponding to unfolding of the binding protein) in the data changes upon mutation to the binding protein while the second transition (corresponding to unfolding of the fluorescent proteins) begins at ~ 90 °C for all variants. The second transition is not observed in full as the upper temperature limit for the experiment is 90 °C.
